# Multi-omics profiling of mouse polycystic kidney disease progression at a single cell resolution

**DOI:** 10.1101/2024.05.27.595830

**Authors:** Yoshiharu Muto, Yasuhiro Yoshimura, Haojia Wu, Monica Chang-Panesso, Nicolas Ledru, Owen M. Woodward, Patricia Outeda, Tao Cheng, Moe R. Mahjoub, Terry J. Watnick, Benjamin D. Humphreys

## Abstract

Autosomal dominant polycystic kidney disease (ADPKD) is the most common hereditary kidney disease and causes significant morbidity, ultimately leading to end-stage kidney disease. PKD pathogenesis is characterized by complex and dynamic alterations in multiple cell types during disease progression, hampering a deeper understanding of disease mechanism and the development of therapeutic approaches. Here, we generate a single nucleus multimodal atlas of an orthologous mouse PKD model at early, mid and late timepoints, consisting of 125,434 single-nucleus transcriptomic and epigenetic multiomes. We catalogue differentially expressed genes and activated epigenetic regions in each cell type during PKD progression, characterizing cell-type-specific responses to *Pkd1* deletion. We describe heterogeneous, atypical collecting duct cells as well as proximal tubular cells that constitute cyst epithelia in PKD. The transcriptional regulation of the cyst lining cell marker GPRC5A is conserved between mouse and human PKD cystic epithelia, suggesting shared gene regulatory pathways. Our single nucleus multiomic analysis of mouse PKD provides a foundation to understand the earliest changes molecular deregulation in a mouse model of PKD at a single-cell resolution.

## Introduction

Autosomal dominant polycystic kidney disease (ADPKD) is a monogenic kidney disease affecting approximately 0.1∼0.2% of the population worldwide(1). The cysts in ADPKD kidneys are induced by a mutation of the polycystin-1 (*PKD1*) or polycystin-2 (*PKD2*) gene, and following deregulation of numerous signaling pathways associated with cAMP response, mammalian target of rapamycin complex (mTORC), WNT and Hippo pathway, among others (1, 2). In addition to disturbed signaling pathways in cystic epithelia, there is also metabolic deregulation of glycolysis, oxidative phosphorylation and lipid metabolism(3). Subsequently, growing cysts compress and injure the adjacent kidney parenchyma and tubular structures, leading to chronic kidney disease (CKD) and ultimately to end-stage kidney disease (ESKD). The vasopressin receptor antagonist tolvaptan slows cyst growth by decreasing cAMP signaling, although this therapy is associated with polyuria, limiting its wide use (4). Therefore, better understanding of disease mechanisms and development of new therapeutic approaches to ADPKD is of paramount importance. However, the dynamic cellular and molecular complexities in ADPKD progression as well as extremely limited access to early stage human ADPKD samples have hampered a deeper understanding of cellular and molecular mechanisms.

Recent evolution in single cell analysis have advanced our knowledge about cellular heterogeneity in healthy and diseased kidneys (5–9). We and others have generated multimodal single cell atlases of human and mouse kidneys, describing previously unrecognized cellular heterogeneity that revealed a novel proinflammatory, pro-fibrotic proximal tubular cell type — failed-repair proximal tubular cells (FR-PTC) (8–12). We have applied our single nucleus multiomics approach to the human ADPKD kidneys, and characterized ADPKD-specific cell states and their molecular signatures (13). We have identified GPRC5A as a novel cyst lining cell marker and its transcriptional deregulation driven by ADPKD-specific epigenetic remodeling. We also revealed cellular heterogeneity of failed-repair proximal tubular cells expanded in human ADPKD kidneys, which may promote PKD progression by modifying the microenvironment (13). Nevertheless, our previous study was limited by the end-stage nature of the ADPKD samples. The cell-type-specific molecular signatures during early stages of PKD progression has remained elusive.

To fill this knowledge gap, we have performed simultaneous snRNA-seq and snATAC-seq on a mouse polycystic kidney disease model over time following *Pkd1* ablation, to comprehensively characterize the cell states and their dynamics during PKD progression. This approach has allowed us to describe the cell-type-specific molecular signatures at an early and middle stage of PKD progression. The atypical collecting duct principal cells as well as failed-repair proximal tubular cells were identified in the PKD mouse kidneys. Comparative analysis of human ADPKD and mouse PKD model atlas sheds light on GPRC5A as a shared cyst lining cell marker, with its transcriptional regulation has been conserved between human and mouse PKD. Our analysis also underscores cell-type-specificity of a molecular response to *Pkd1* deletion during PKD progression, rather than a shared response to *Pkd1* deficiency across tubular compartments. This single-nucleus multiomics atlas as well as our findings through the analysis will be the foundation to better understand the mechanism of PKD and develop novel therapeutic approaches.

## Results

### Single nucleus multimodal profiling of slow-onset PKD model mouse kidneys

To comprehensively understand the cell-type-specific molecular mechanism of PKD progression, we applied single-nucleus multiomics to an orthologous mouse model of ADPKD — the *Pkd1*^*fl/fl*^; *Pax8*^*rtTA*^; *TetO*^*Cre*^ mouse that utilizes the *Pax8* promoter to drive expression of a reverse tetracycline-dependent transactivator (rtTA) in all renal tubular compartments including proximal and distal tubules as well as collecting ducts(14, 15). We induced *Pkd1* deletion in the tubular cells of these male mice from postnatal day 27 (P27) to 57 (P57) with doxycycline injection. The littermate male controls lacked either the *Pax8*^*rtTA*^ (P66) or the *TetO*^*Cre*^ module (P100, P130, Table S1) but were treated with doxycycline in the same manner. The kidneys were harvested and snap-frozen at postnatal day 66 (P66), 100 (P100) or 130 (P130) (Fig. 1A, Table S1). The PKD mouse kidneys demonstrated a gradual increase in the ratio of kidney weight to body weight (Fig. 1A, B, Table S1), along the progression of PKD (Fig. S1). Immunostaining of tubular lineage markers indicated that the cysts in this PKD model kidneys derived from multiple tubular compartments including lotus tetragonolobus lectin (LTL)+ proximal tubules, uromodulin (UMOD)+ thick ascending limbs of Henle loop and lectin Dolichos biflorus agglutinin (DBA)+ collecting ducts (Fig. 1C). Of note, DBA+ collecting ducts already mildly dilated as early as at P66 in corticomedullary junction (Fig. 1C), suggesting ongoing cystic transformation of these cells. In contrast, tubules in the cortex did not show apparent morphologic changes at P66 (Fig. 1C) except occasional small cysts (Fig. S1). Both cortical and corticomedullary regions demonstrated numerous cysts at P100, and the cystic lesions were most prominent at P130 (Fig. 1B, C, Fig. S1). These findings suggest that our multiomics atlas includes the major PKD stages from a pre-cystic to late cystic stages. We generated and sequenced multimodal single-nucleus libraries on a total of 16 mouse kidney samples (8 PKD and 8 control mouse kidneys) along a time course following *Pkd1* deletion (Fig. 1A).

**Figure 1.**
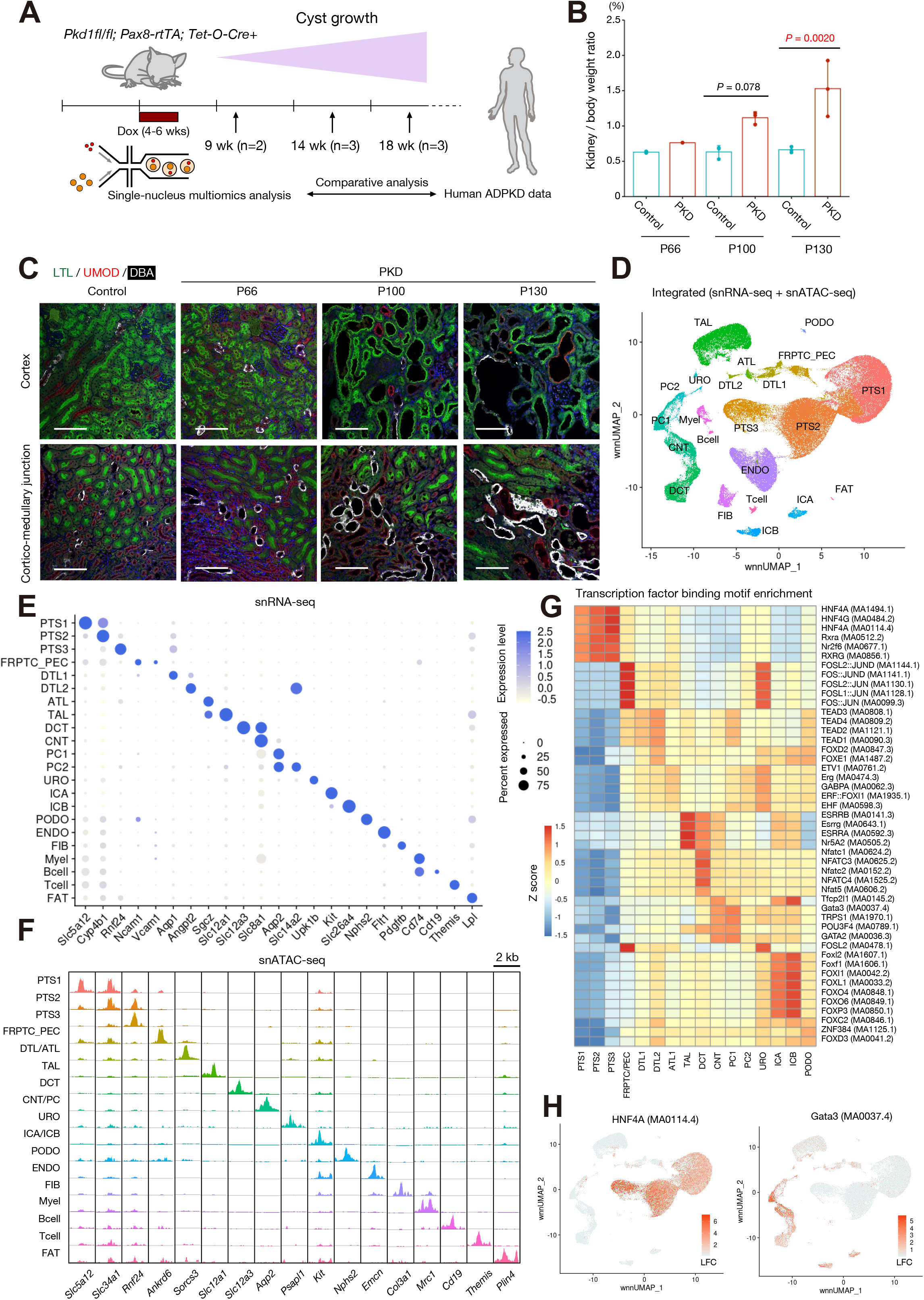
Single nucleus multiomics profiling for mouse PKD kidneys. **(A)** Overview of experimental strategy. Single nucleus multiomics atlas was generated from PKD model mouse kidneys and littermate control kidneys along a time course (three time points; postnatal day 66 (P66), 100 (P100) or 130 (P130) after *Pkd1* deletion (n = 2-3 pairs for each time point). (**B**) The kidney to body weight ratio of the PKD and control mice that we have analyzed by multiomics analysis. Bar graphs represent the mean and error bars are the s.d. One-way ANOVA with post hoc Turkey test. (**C**) Representative immunofluorescence images of lotus tetragonolobus lectin (LTL, green), uromodulin (UMOD, red) and lectin Dolichos biflorus agglutinin (DBA, white) in the cortex (upper) or corticomedullary junction (lower) in the kidneys (control [P100], PKD [P66, P100 and P130]). Scale bar indicates 100 μm. (**D**) UMAP plot of the integrated single nucleus multiomics dataset with weighted nearest neighbor (wnn) clustering. Clusters were annotated by lineage marker gene expression. PTS1/S2/S3, proximal tubule S1/S2/S3 segments; FRPTC_PEC, failed-repair proximal tubular cells and parietal epithelial cells; DTL1/DTL2/ATL, descending thin limb 1/2 and ascending thin limb of Henle’s loop; TAL, thick ascending limb of Henle’s loop; DCT, distal convoluted tubule; CNT, connecting tubule; PC1/2, principle cells 1/2; URO, uroepithelial cells; ICA, Type A intercalated cells; ICB, Type B intercalated cells; PODO, podocytes; ENDO, endothelial cells; FIB, fibroblasts; Myel, myeoloid cells; FAT, adipocyte. (**E**) Dot plot showing gene expression patterns of cluster-enriched markers for the integrated dataset. The diameter of the dot corresponds to the proportion of cells expressing the indicated gene and the density of the dot corresponds to average expression relative to all cell types. (**F**) Fragment coverage (frequency of Tn5 insertion) around the differentially accessible regions (DAR) around each cell type at lineage marker gene transcription start sites. Scale bar indicates 2 kbp. (**G**) Heatmap showing averaged enrichment for the most enriched transcription factor binding motifs in each cell type. (**H**) UMAP plot showing chromVAR motif enrichment scores in the dataset for HNF4A (MA0114.4, left) and GATA3 (MA0037.4, right). The color scale represents a normalized log-fold-change (LFC).

After sequencing (Table S2), batch quality control (QC) filtering and preprocessing of sequencing data (Fig. S2), the individual multiomics datasets were integrated with Seurat (16). We obtained a total of 125,434 nuclei (59,089 nuclei from PKD and 66,345 nuclei from control). Both control and PKD data show similar numbers of unique genes and transcripts per nucleus (Fig. S3), indicating successful generation of high-quality dataset from the samples including those with advanced PKD (Fig. 1B). Following dimensional reduction on both transcriptomic and epigenomic data with batch effect correction with Harmony (17), a Weighted Nearest Neighbor (WNN) graph was calculated and visualized on UMAP space (Fig 1D). The cell types were annotated by marker gene expression (Fig. 1E). The lineage marker expression were largely maintained during progression of PKD (Fig. S4), consistent with our previous finding that the cell type marker gene expression was preserved in human advanced ADPKD kidneys (13). All major cell types were identified in both control and PKD mouse kidneys (Fig. S5, Table S3) but a cluster representing failed-repair proximal tubular cells (FR-PTC) gradually expanded in PKD along the time course (Fig. S5, Table S3). Although immune cell infiltration as well as interstitial fibrosis have been observed in advanced PKD, the increase in the numbers of those cell types were not observed in our dataset, presumably due to dissociation bias(18).

Next, we catalogued differentially expressed genes in PKD for each cell type at each time point. Surprisingly, there were only a few differentially regulated genes shared among tubular cell types (PT, TAL, PC1/2) in PKD (P66, *Kap* and *Fkbp5*; P100, *Nox4* and *Gm42418*; P130, no shared genes) (Fig. S6A). This finding suggests that the molecular response to *Pkd1* deletion is cell-type-specific, although the consequences — cystogenesis and cyst growth — are shared across tubular compartments. This observation was in line with the possibility that the mechanism of cystogenesis and cyst growth may be cell-type-specific, underscoring the importance of single cell analysis to dissect PKD mechanisms.

### Cell-type-specific chromatin accessibility and motif availability in PKD

The cell types annotated through marker gene expressions (Fig.1E) demonstrated consistent cell-type-specific chromatin accessibility to marker genes (Fig. 1F). Dimensional reduction and projection of nuclei on UMAP with snATAC-seq peaks also indicates that clustering based on chromatin accessibility is largely in line with that with transcriptomic data (Fig. S7) or integrated approach (Fig. 1D). In agreement with the transcriptional data, there was only one shared differentially accessible region (DAR) among PT, TAL and PC in PKD at P66 (chr2:24478513-24480280, ∼3 kb 5’-distal to *Pax8* gene) and no shared DAR at P100 or P130 (Fig. S6B). The shared DAR in the *Pax8* locus at P66 PKD simply reflects a lack of the *Pax8*-rtTA transgene in controls. This finding emphasizes the cell-type-specificity of epigenetic changes induced by *Pkd1* deletion. Next, we performed transcription factor binding motif enrichment analysis on accessible chromatin regions at a single nucleus level (Fig. 1G, H) with ChromVar(19). The transcription factors enriched in certain cell types were often associated with those lineage specifications (Fig. 1H) (8).

Next, we compared the enriched transcription factor binding motifs between PKD and control kidneys. One of the most enriched transcription factor motifs in late stage PKD (P130) in multiple tubular cell types (PT, TAL, PC1, PC2) was Transcription factor EB (TFEB), a master regulator of lysosomal function and autophagy (Fig. S8A). Nuclear localization and noncanonical activation of TFEB in cystic epithelia was previously shown in the mouse and human polycystic kidneys(20). TFEB was also found to drive mTOR hyperactivation and renal cell carcinoma in tuberous sclerosis complex(21). Constitutive activation of TFEB and its family transcription factor TFE3 promote cell proliferation in several types of cancers in addition to renal cell carcinoma(22). These lines of evidence may implicate TFEB in aberrant growth of cyst epithelia in PKD. AP-1 transcription factor binding motifs were also enriched in tubular cell types in PKD (Fig. S8B), presumably reflecting a cohort of proinflammatory signaling pathways activated in advanced PKD (23). These findings collectively suggest that alteration in the chromatin accessibility landscape was associated with differential transcription factor availability during PKD progression, potentially promoting activation of cystic epithelia and inflammation in PKD.

### Deregulation of molecular pathways in PKD collecting duct cells

We have previously characterized cellular heterogeneity of collecting duct cells in human ADPKD, revealing activation of inflammatory pathways as well as metabolic deregulation in cystic cell types (13). The mouse PKD kidneys also have numerous DBA+ distal nephron cysts (Fig. 1C). To characterize the mouse PKD distal nephron cells, we performed sub-clustering analysis for the distal nephron cell types; connecting tubules (CNT), principal cells (PC1 and PC2) and uroepithelial cells (URO) in the whole dataset (Fig. 1D), identifying 8 subpopulations (Fig. 2A). These subtypes express unique marker genes (Fig. 2B). CNT subtypes express *Ptprd, Slc8a1* and *Calb1* as shared marker genes. The cortical principal cells (cPC1/2) and medullary principal cells (mPC1/2) express *Apq2*, which encodes aquaporin2 (AQP2). All these subtypes were detected in both PKD and control at each time point (Fig. S9). Immunostaining of CALB1 and AQP2 proteins in the late stage PKD kidneys revealed that most of cystic CALB1+ cells were also AQP2+/DBA+, and that CALB1+ tubules lacking AQP2 expression rarely contributed to cyst formation (Fig. 2C, Fig. S10), suggesting that the majority of DBA+ cysts observed in PKD kidneys were originated from collecting ducts. To compare the mouse PKD principal cells with human ADPKD principal cells (13), we predict the corresponding subtypes of mouse PKD distal nephron in the human ADPKD distal nephron cells. Label transfer of subtype annotations from mouse PKD to human ADPKD indicates that the majority of human ADPKD cystic collecting duct cells (PKD-CDC)1 were related to the mPC2 population, and human PKD-CDC2 cells were associated with cPC1 cluster (Fig. 2D) (13). The gene expression signature of PKD-CDC1/2 in human ADPKD was associated with inflammation, hypoxia and cellular senescence, with up-regulation of cyst lining cell marker GPRC5A (13). The cell-type prediction suggests that the cellular heterogeneity of collecting duct cells are conserved between human and mice.

**Figure 2.**
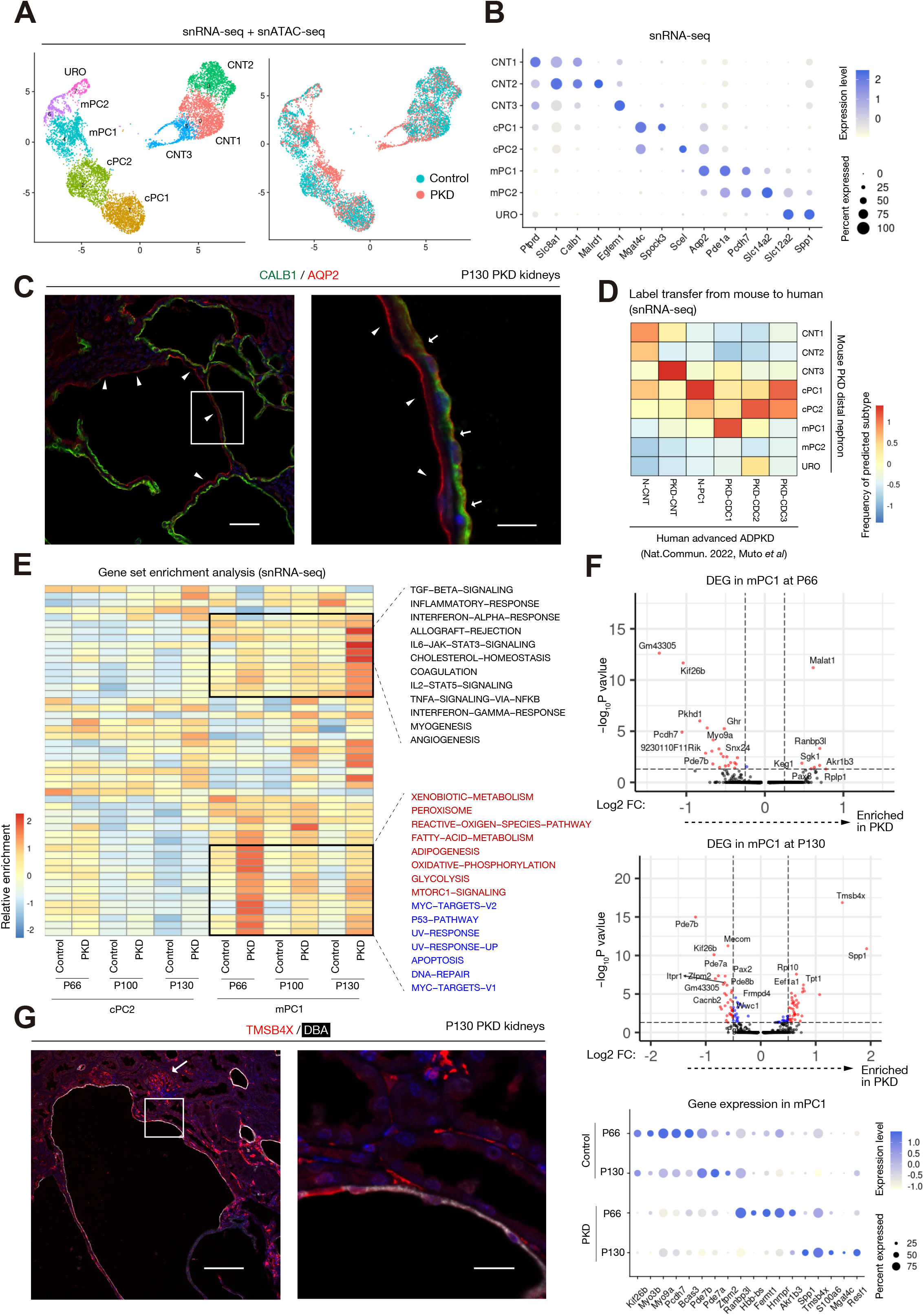
Heterogeneity of collecting duct principal cells in mouse PKD. (**A**) Sub-clustering of distal nephron clusters (CNT, PC1, PC2 and URO) on the UMAP plot. cPC, cortical principal cells; mPC, medullary principal cells. (**B**) Dot plot showing expressions of the genes enriched in each of the subtypes. The diameter of the dot corresponds to the proportion of cells expressing the indicated gene and the density of the dot corresponds to average expression. (**C**) Representative immunofluorescence images of CALB1 (green) and AQP2 (red) in the PKD kidneys at P130. Arrowheads indicate AQP2+ cyst lining. Arrows mark AQP2+/CALB1+ cyst lining. Scale bar indicates 100 μm (left) or 12.5 μm (right). (**D**) Distal nephron subtypes in the human advanced ADPKD data are label-transferred from those in the mouse PKD data, and the frequencies of predicted mouse subtypes in each human subtype are shown on the heatmap. (**E**) Heatmap showing enrichment of gene expressions of the hallmark gene sets among cPC2 and mPC1 clusters at each time point. The pathways associated DNA damage response are in blue characters and those associated with metabolic regulation are in red characters (**F**) Volcano plot showing differentially expressed genes in mPC1 of PKD mice compared to that of control at P66 (upper) or P130 (middle). The x-axis represents the log fold change, and the y-axis represents the *p* value with corresponding expressions. Dot plot showing differential gene expressions in mPC1 between PKD and control at P66 (upper) or P130 (middle). Dot plot (low) showing expressions of the representative genes in mPC1 at P66 and P130. The diameter of the dot corresponds to the proportion of cells expressing the indicated gene and the density of the dot corresponds to average expression. (**G**) Representative immunofluorescence images of DBA (green) and TMSB4X (red) in the PKD kidneys at P130. An arrow indicates a glomerulus. Scale bar indicates 100 μm (left) or 15 μm (right).

To better understand the molecular signature of mPC2 and cPC1 subtypes in mouse PKD, we applied gene set enrichment analysis on cPC2 and mPC1 subtypes (Fig. 2E). Interestingly, the differentially regulated signaling pathways in these subpopulations were dynamically changed during PKD progression. We observed activation of a cohort of genes associated DNA damage response (Fig. 2E, pathways in blue characters) and metabolic alterations (Fig. 2E, pathways in red characters) at P66 in mPC1, implicating activation of these pathways in early morphological changes in the collecting duct (Fig. 1C). In contrast, gene expression associated with inflammatory pathways are up-regulated in mPC2 at the late stage PKD samples (P130), presumably reflecting inflammation secondary to tissue destruction by large cysts as well as recruitment of immune cells (Fig. 2E). This observation in late-stage PKD was also in line with activation of inflammatory pathways in PKD-CDC1/2 in human advanced ADPKD (13). Consistent with dynamic alterations of gene set enrichment in mPC1, the differentially expressed genes in mPC1 (Fig. 2F) as well as cPC2(Fig. S11) are disease-stage-specific (Fig. 2F). Among the differentially upregulated genes in the late stage mPC2, the most up-regulated gene was *Tmsb4x* that encodes thymosin beta 4 (TMSB4X) protein, which is an actin sequestering protein playing a role in regulation of cytoskeleton(24, 25). TMSB4X has been shown to be beneficial in diverse pathological conditions including myocardial infarction, neuronal damage and diabetic nephropathy (24, 25). In contrast, TMSB4X has been found to promote tumor progression and to be associated with poor prognosis in various cancers(24). Immunostaining analysis identified expression of TMSB4X in DBA+ cysts (Fig.2G), implicating TMSB4X in cyst growth in advanced PKD. *Tmsb4x* is broadly up-regulated in multiple cell types including parenchymal cell types including endothelial cells and immune cells (Fig. S12), in line with interstitial staining of TMSB4X in addition to cyst lining cells (Fig. 2G). The relatively high level of *Tmsb4x* expression in podocytes was also consistent with previous study (Fig. 2G, Fig. S12) (25). These findings collectively suggest that heterogeneous principal cell states show dynamic alterations in their molecular signatures and signaling pathways during PKD progression.

### Heterogeneous failed-repair proximal tubular cells in cystic epithelia of mouse PKD

We and others have described *Vcam1*-expressing failed-repair proximal tubular cells (FR-PTC) after kidney injury in mice (10–12, 26) and human(8, 9, 13, 27). This subpopulation is characterized by pro-inflammatory, pro-fibrotic gene expression signature, implicated in progression of kidney diseases(13, 27). Of note, the frequency of FR-PTC in advanced ADPKD was markedly increased, replacing normal PTC(13), suggestive of its significant role in ADPKD progression(13). Furthermore, we also revealed the heterogeneity of FR-PTC in human ADPKD kidneys.

We also observed expansion of FR-PTC cluster during mouse PKD progression (Fig. 1D, Fig. S5, Table S3). To understand the heterogeneity of FR-PTC in PKD mouse kidneys, we performed sub-clustering analysis on the FRPTC_PEC cluster (Fig. 1D), identifying 4 epithelial subtypes besides PEC (Fig. 3A, B). Cluster 0 express normal PT lineage markers, although they are down-regulated compared to normal epithelia (Fig. 3B), suggesting that they may be a transitional state (Transitional PTC). FR-PTC1 highly express previously described FR-PTC marker genes; *Havcr1* and *Vcam1* (Fig. 3B, C). In agreement with this observation, *Vcam1* gene promoter is more accessible in FR-PTC1 (Fig. 3C) compared to other PT lineages. Immunostaining analysis of PKD kidneys at a cystic stage revealed accumulation of VCAM1+ cells in the cyst lining and non-cystic, atrophic tubules throughout the cortex (Fig. 3D). Interestingly, VCAM1 protein expression was mutually exclusive to LTL staining even in the same cysts (Fig. 3D), suggesting that VCAM1 is a marker for dedifferentiation of the proximal cystic epithelia. Furthermore, cysts with VCAM1+ epithelia often acquire epithelial lining lacking both LTL and VCAM1 (Fig. 3D). Consistent with this observation, FR-PTC2/3 subsets demonstrated much lower levels of *Vcam1* expression compared to FRPTC1 (Fig. 3B, C). FR-PTC2 and FR-PTC3 demonstrated similar gene expression signatures (Fig. 3B). However, the general chromatin accessibility of FR-PTC3 was limited compared to other subtypes (Fig. S13). The poor chromatin accessibility in FR-PTC3 may be due to extensive chromatin condensation.

**Figure 3.**
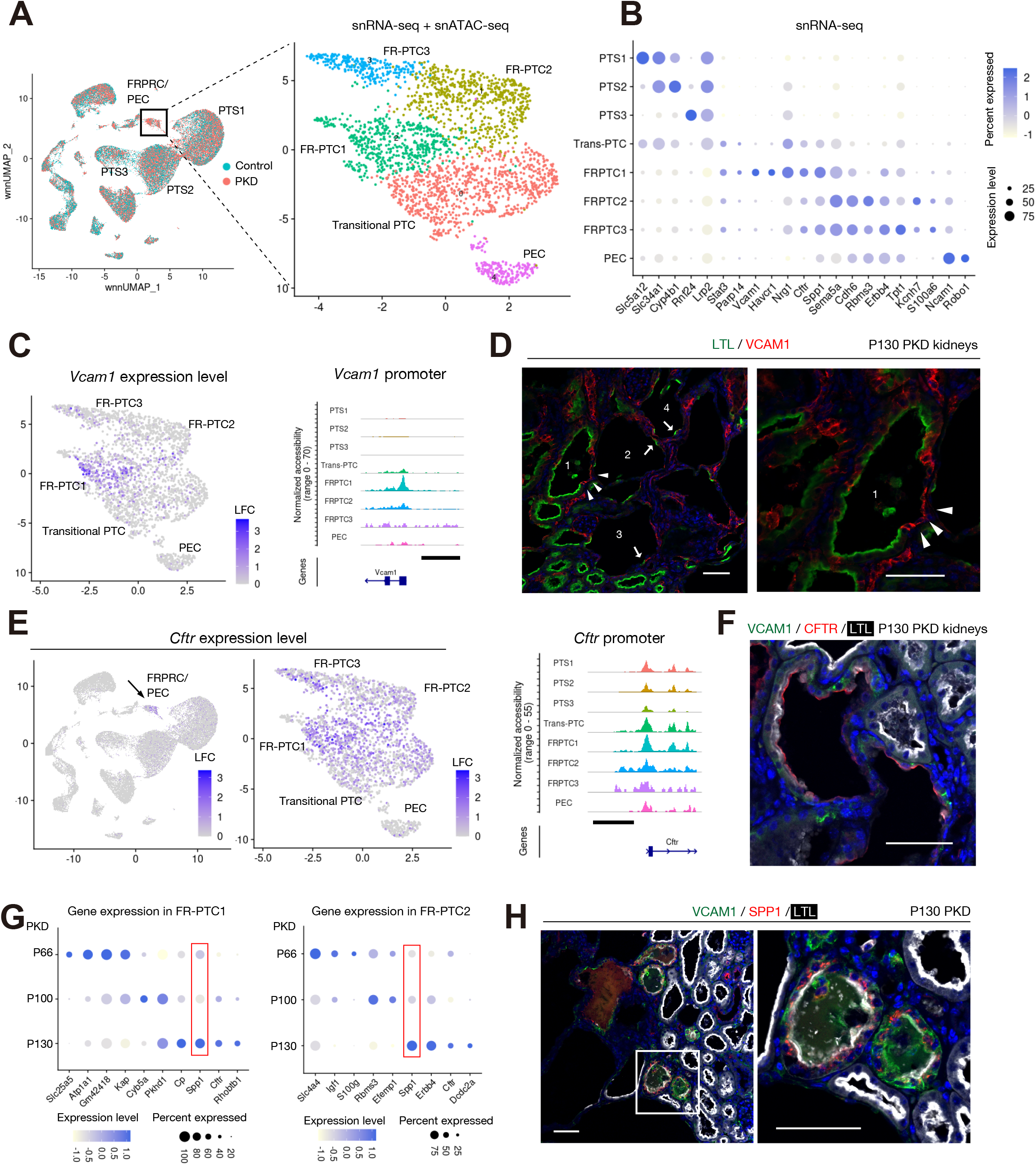
Heterogeneity of failed-repair proximal tubular cells in mouse PKD. (**A**) Sub-clustering of FRPTC_PEC cluster on the UMAP plot. (**B**) Dot plot showing expressions of the genes enriched in each of the subtypes. The diameter of the dot corresponds to the proportion of cells expressing the indicated gene and the density of the dot corresponds to average expression. (**C**) UMAP plot showing a gene expression level of *Vcam1* (left) and coverage plot showing accessible regions around *Vcam*1 promoter in each subtype was also shown (right). The color scale represents a normalized log-fold-change (LFC, left). The scale bar indicates 2 kbp (right). (**D**) Representative immunofluorescence images of LTL (green) and VCAM1 (red) in the PKD kidneys at P130. Arrowheads indicate VCAM1+ FR-PTCs replacing LTL+ PTC in the cyst1. Arrows mark LTL+ cyst lining in the cysts2-4, which have acquired LTL-VCAM1-epithelial lining. Scale bar indicates 50 μm. **(E**) UMAP plot showing a gene expression level of *Cftr* (left) and coverage plot showing accessible regions around *Cftr* promoter in each subtype (right). The color scale represents a normalized LFC. The scale bar indicates 2 kbp (right). (**F**) Representative immunofluorescence image of VCAM1 (green), CFTR (red) and LTL (white) in the PKD kidneys at P130. Scale bar indicates 50 μm. (**G**) Dot plot showing expressions of the differentially expressed genes at different time points in FRPTC1 (left) or FRPTC2 (right). The diameter of the dot corresponds to the proportion of cells expressing the indicated gene and the density of the dot corresponds to average expression. (**H**) Representative immunofluorescence images of VCAM1 (green), SPP1 (red) and LTL (white) in the PKD kidneys at P130. Scale bar indicates 50 μm.

Despite a lack of *Vcam1* expression, FR-PTC2/3 subtypes shared the expression of *Cftr, Sema5a* and *Spp1* with FR-PTC1 (Fig. 3B). Interestingly, *Cftr* expression was specifically up-regulated in FR-PTCs in PKD (Fig. 3E, Fig. S14A), while its expression was minimal in the FR-PTC in mouse kidneys with ischemia-reperfusion injury (Fig. S15A)(10), suggestive of a PKD-specific mechanism to up-regulate *Cftr* gene expression. *Cftr* gene encodes cystic fibrosis transmembrane conductance regulator (CFTR) protein, which has been implicated in fluid accumulation in cysts (28). Immunostaining analysis suggests an apical CFTR protein localization in both VCAM1+ and LTL+ cysts in mouse PKD kidneys (Fig. 3F), although we also observed PT-derived cysts lacking CFTR expression, suggesting additional translational or post-translational regulation for the CFTR protein expression or localization, which may be designated by microenvironmental cues that activate cellular cAMP signaling (29).

The gene expression pattern in each FRPTC subtype also changed dynamically during PKD progression (Fig. 3G), which was also observed in collecting duct subpopulations (Fig. 2). The genes with dynamically changed expression during PKD progression include *Spp1* and *Cftr* in both FRPTC1 and FRPTC2 (Fig. 3G). *Spp1* encodes a pleiotropic glycoprotein; osteopontin (SPP1), which has been shown to be involved in inflammation, angiogenesis and apoptosis in pathologic conditions including ADPKD(30). We observed SPP1 protein localized in cystic epithelia with PT lineage marker expression (Fig. 3H). *Spp1* expression was widely detected and up-regulated in other tubular and non-tubular cell types in PKD kidneys (Fig. S14B), consistent with a recent line of evidence suggesting that *Spp1* expression in the pericystic endothelium in PKD(31). SPP1 expression in cyst epithelia (Fig. 3B, H) may alter surrounding microenvironment, potentially promoting PKD progression.

Next, we asked whether the heterogeneity of FR-PTC in mouse PKD is conserved in human ADPKD. Label transfer of these FR-PTC subtype annotations from mouse PKD (Fig. 4A) to human ADPKD FR-PTC predicted that the gene expression signatures of mouse FR-PTC2 was closest to that of human ADPKD PT3 subtype (Fig. 4A), which was characterized by a low expression level of *VCAM1(13)*. This finding indicates conservation of cellular heterogeneity in FR-PTC between human and mouse PKD. Together, these observations implicate heterogeneous FR-PTC subtypes in PKD progression as components of cystic epithelia. Furthermore, the gene expression signature of each FR-PTC subtype is dynamically altered during PKD progression (Fig. 3G).

**Figure 4.**
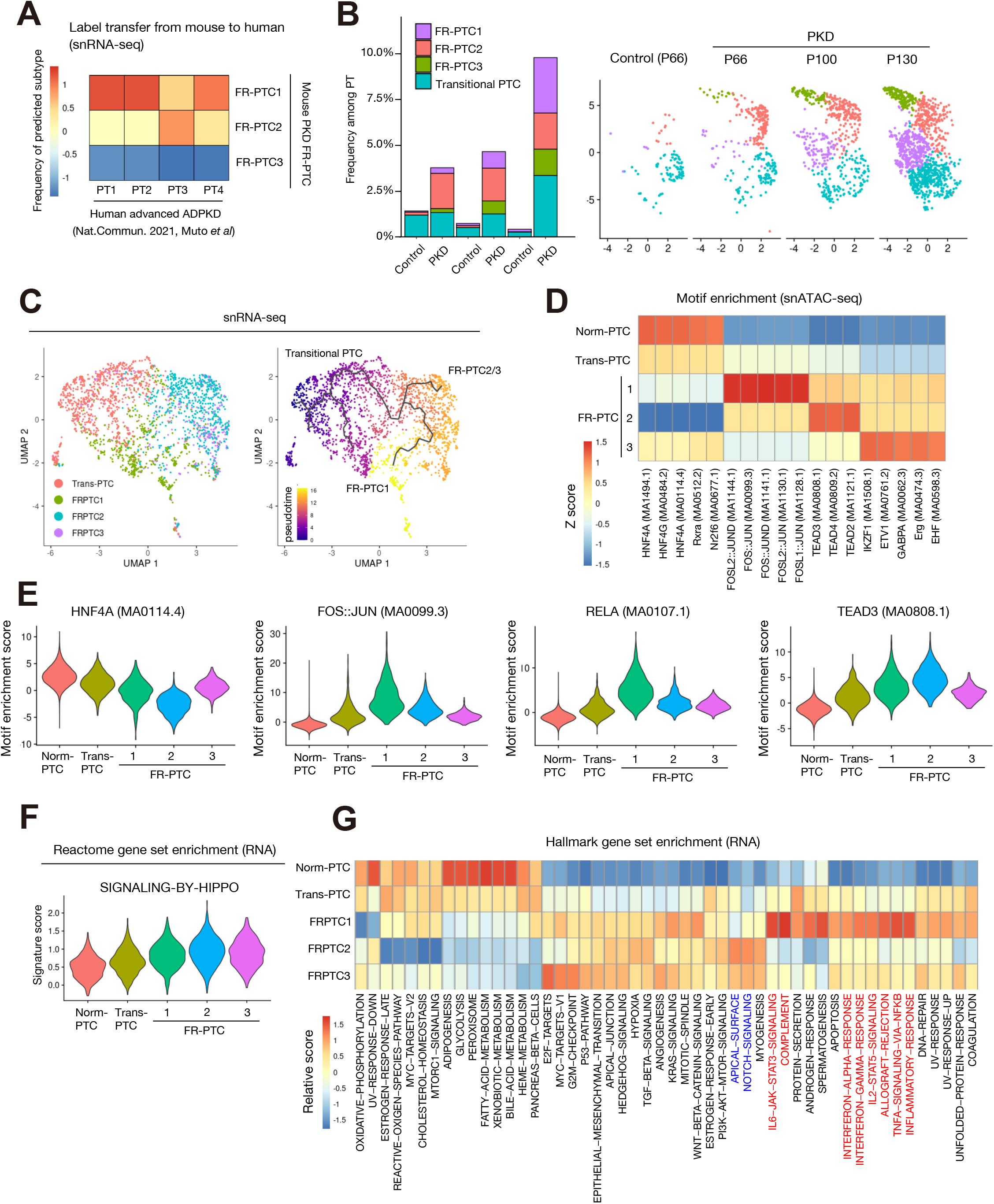
Differentially activated molecular signaling pathways among PT subtypes in PKD. (**A**) Proximal tubular cell subtypes of human advanced ADPKD data are label-transferred from FR-PTC subtypes in mouse PKD data, and the frequencies of predicted mouse subtypes in each human subtype are shown on the heatmap. (**B**) Frequency of each PT subtype among whole PTC for each time point in PKD or control data (left). UMAP plot showing FRPTC subtypes at each time point (right). (**C**) Pseudotemporal trajectories to model differentiation from transitional PT to FR-PTC subtypes, constructed with snRNA-seq. Colored by the subtypes (left) or pseudotime (right). (**D**) Heatmap showing relative transcription factor binding motif enrichment among PT subtypes in snATAC-seq. Most enriched transcription binding motifs in each subtype are shown. (**E**) Violin plots displaying relative motif enrichment (chromVAR score) among PTC for HNF4A (MA0114.4), JUN::FOS (MA0099.3), RELA (MA0107.1) or TEAD3 (MA0808.1) (**F**) Violin plots displaying relative gene set enrichment among PTC for REACTOME-SIGNALING-BY-HIPPO. (**G**) Heatmap showing relative enrichment of hallmark gene sets among PT subtypes in snRNA-seq. The pathways related to cell polarity and notch signaling were in blue characters, and the inflammatory pathways are in red characters.

### Atypical failed-repair proximal tubular cells emerging after PKD1 inactivation

The frequency of FRPTC1 subtype increased during PKD progression (Fig. 4B), reflecting accumulation of VCAM1+ cells in PKD kidneys (Fig. 3D). In contrast, the frequency of FRPTC2 was relatively steady during PKD progression, suggesting that FRPTC2 may be a reversible cell state. FR-PTC2 specifically up-regulated the expression of *Cdh6* encoding cadherin 6 (CDH6) (Fig. 3A), which was recently shown to be a marker of a distinct, regenerative cell state following epithelial injury and deregulation of cell polarity (32). FR-PTC2 may be the dedifferentiated cell state directly induced by *Pkd1* deletion. Pseudotemporal analysis suggested that the trajectory from transitional PTC to FRPTC1 was independent from that to FRPTC2 (Fig. 4C), indicating that FRPTC1 and FRPTC2 may be separately dedifferentiated from normal epithelia.

Next, we performed transcription factor binding motif enrichment analysis to understand the epigenetic mechanism driving the heterogeneity of FRPTCs (Fig. 4D). FRPTC1 enriched AP-1 transcription factor (Fig. 4D, E) as well as NF-kB family transcription factor binding motifs (Fig. 4F), consistent with previous findings that *Vcam1*+ FR-PTC demonstrates an increase in AP-1 and RELA binding motif availability(8, 12, 33). In contrast, the FRPTC2 subtype showed motif enrichment of TEAD family transcription factors (Fig. 4D, E), through which Hippo signaling pathway regulates expression of the downstream target genes. Hippo pathway has been implicated in regulation of proliferation during development and pathologic condition including various cancers(34) and ADPKD (35). The most differentially up-regulated gene in FRPTC2 was *Rbms3* encoding an RNA-binding protein (Fig. 3B). *Rbms3* was found to be one of the common Hippo pathway target (34), in agreement with Hippo pathway deregulation in this subtype. Furthermore, gene set enrichment analysis indicated up-regulation of genes related to Hippo pathway in FR-PTC, especially FRPTC2 (Fig. 4F). Gene set enrichment analysis for hallmark gene sets suggested enrichment of genes related to cell polarity (apical surface) as well as notch signaling in FRPTC2 (pathways in blue characters in Fig. 4G). Both of these pathways have been implicated in deregulation of Hippo pathway(36, 37). In contrast, FRPTC1 enriched inflammatory pathway genes (pathways in red characters in Fig. 4G).

Collectively, our analysis suggests each of FRPTC subtypes separately dedifferentiated from normal epithelia by *Pkd1* deletion, with unique activation of transcription factors and downstream molecular signaling. These FRPTC subtypes may have unique roles in PKD progression.

### GPRC5A as a cyst lining cell marker for both human and mouse PKD

We have previously identified GPRC5A as a marker for cyst-lining cells in human advanced ADPKD samples(13). GPRC5A is a G-protein-coupled receptor which is deregulated in many forms of cancer(38). Recently aromatic monoamines were found to bind GPRC5A and stimulate β-arrestin recruitment (39), although the physiological ligands in vivo as well as its biological function has been elusive. *Gprc5a* expression was detected in many cell types although its expression was most abundant in FRPTC/PEC, DTL as well as some distal nephron cell types (PC2/URO) in our dataset (Fig. 5A). In the distal nephron sub-clustering, *Gprc5a* expression was mainly detected in cPC2, mPC1/2 as well as URO (Fig. 5B). GPRC5A protein expression was often observed in DBA+ collecting duct cyst lining cells (Fig. 5C). *Gprc5a* expression was also widely detected in FRPTC subtypes, while it was hardly detected in transitional PTC (Fig. 5D), consistent with the observation that FRPTC composes cystic epithelia in the PKD kidneys (Fig. 3). Immunostaining analysis revealed GPRC5A protein was often detected in proximal tubular cysts (Fig. 5E). Similar to VCAM1, GPRC5A+ epithelia lost LTL staining, indicating GPRC5A is a marker for dedifferentiation in PT-derived cyst epithelia. In agreement with this notion, GPRC5A expression was often colocalized with VCAM1 expression (Fig. 5F). GPRC5A expression was not observed in VCAM1+ non-cystic, atrophic tubules, suggesting that GPRC5A is a cyst cell marker but not FR-PTC marker (Fig. 5F). In agreement with this observation, *Gprc5a* up-regulation was not observed in FR-PTC in the mouse kidneys with IRI (Fig. S15B). These findings collectively indicated that GPRC5A is a conserved cyst lining cell marker between human and mouse PKD, and that it marks the cyst lining cells regardless of their original lineages.

**Figure 5.**
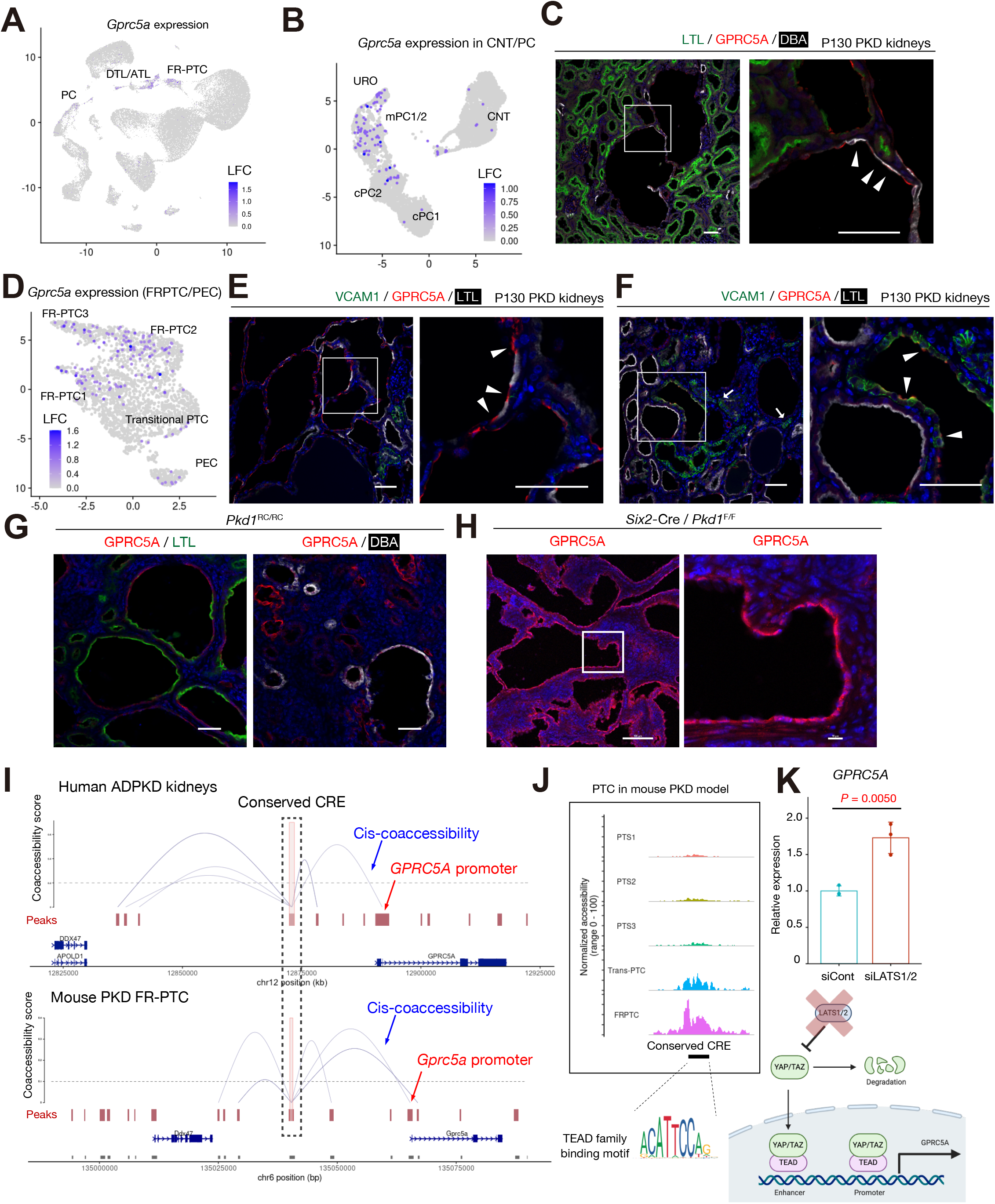
GPRC5A as a shared cyst lining cell marker for mouse and human PKD. (**A, B**) UMAP plot showing a gene expression level of *Gprc5a* in the whole dataset (A) or distal nephron subclustering (B). The color scale represents a normalized log-fold-change (LFC). (**C**) Representative immunofluorescence images of LTL (green), GPRC5A (red) and DBA (white) in the PKD kidneys at P130. Arrowheads indicate GPRC5A+DBA+ cyst lining. Scale bar indicates 50 μm. (**D**) UMAP plot showing a gene expression level of *Gprc5a* in the FR-PTC sub-clustering. The color scale represents a normalized LFC. (**E, F**) Representative immunofluorescence images of VCAM1(green), GPRC5A (red) and LTL (white) in the PKD kidneys at P130. Arrowheads indicate cyst lining with GPRC5A mutually exclusive with LTL (E) or colocalized with VCAM1 (F). The arrows indicate VCAM1+ atrophic tubules. Scale bar indicates 50 μm. (**G, H**) Representative immunofluorescence images of LTL (green), GPRC5A (red) and DBA (white) in the *Pkd1*^RC/RC^ mouse cystic kidneys at 11 months of age (G) or *Six2*-Cre; *Pkd1*^F/F^ cystic kidneys at P7 (H). Scale bar indicates 50 μm (G) or 100, 10 μm (H). (**I**) Cis-coaccessibility network (CCAN, gray arcs) of a conserved cis-regulatory region (CRE) 5’ distal to *Gprc5a* promoter in the mouse PKD FR-PTC (lower) or human ADPKD (upper) among accessible regions (red boxes) is shown. (**J**) Coverage plot showing accessibility of conserved CRE 5’ distal to *Gprc5a* gene among PT subtypes (upper). The conserved CRE has several TEAD family binding motifs both in human and mice. (**K**) Quantitative PCR for *GPRC5A* expression in primary human PTC with siRNA knockdown of *LATS1* and *LATS2* (upper). n = 3 biological replicates with two-sided Student’s t-test. *LATS1/2* knockdown inhibits Hippo pathway, activating TEAD and subsequently up-regulating *GPRC5A* expression (lower). Bar graphs represent the mean and error bars are the s.d. Student’s t-test. Schematic was created with BioRender.

*Pkd1*^RC/RC^ is another orthogonal ADPKD model mouse with Arg3277Cys (RC) mutation, which is associated with hypomorphic PKD1 (40). The cystic kidneys with this mouse model demonstrated GPRC5A expression in the cyst lining cells originated from proximal tubular cells as well as collecting duct cells (Fig. 5G). Furthermore, the cystic lesions in a rapidly progressive PKD model mouse (*Six2*-Cre; *Pkd1*^F/F^) kidneys (41) also demonstrated localization of GPRC5A in the cyst lining cells (Fig. 5H). These findings confirmed GPRC5A is a common cystic cell marker in disparate mouse models of PKD.

### Conservation of GPRC5A distal enhancer and regulation of GPRC5A through Hippo pathway

We identified a cis-regulatory region (CRE) 5’ distal to the GPRC5A promoter in human ADPKD kidneys and validated it by CRISPR interference(13). The sequence of the CRE was conserved on the 5’ distal region of the *Gprc5a* gene also in the mouse genome (remapping ratio > 0.5, Fig. 5G), and this region was more accessible in FR-PTC compared to normal PT epithelia in the PKD mouse model (Fig. 5I, J). Cis-coaccessibility network (CCAN) modeling in mouse FR-PTC predicted the interaction between this conserved CRE with the *Gprc5a* promoter (Fig. 5I), suggesting that the epigenetic mechanism regulating *Gprc5a* expression is conserved in mouse PKD. We have also shown that the CRE in human ADPKD includes cAMP-responsive element binding protein 1 (CREB1) motif as well as Hippo pathway effector TEAD family transcription factor binding motifs(13). The mouse *Gprc5a* CRE also have TEAD binding motifs (Fig. S16). ChIP-seq of TEAD1 in a human cancer cell lines have shown TEAD1 coupled with YAP bind to the GPRC5A CRE (Fig. S17)(42), suggesting that the CRE may regulate GPRC5A expression through Hippo pathway, in addition to cAMP and retinoic acid signaling (13). LATS1/2 are the kinases essential for Hippo pathway regulation, phosphorylating YAP for subsequent degradation. Deregulation of the Hippo pathway by siRNA knockdown of LATS1/2 induced GPRC5A up-regulation in the primary proximal tubular cells (Fig. 5K), in agreement with the notion that GPRC5A is regulated by TEAD family transcription factors in addition to cAMP and retinoic acid signaling pathways. This finding along with our previous study(13) indicates that GPRC5A is regulated by a cohort of signaling pathways associated with PKD progression, including cAMP, retinoic acid and Hippo pathways. These findings collectively suggest that the epigenetic mechanism of GPRC5A regulation and the gene regulatory network driving PKD progression are largely conserved between human and mouse kidneys.

## Discussion

In this study, we generated a single-nucleus multiomic atlas for a mouse model of PKD along the time course following *Pkd1* inactivation to comprehensively describe the dynamic cell states during PKD progression. This research strategy allows us to dissect cellular heterogeneity and characterize molecular response to PKD1 deficiency in each cell type at each disease stage. Given the recent evidence that PKD is partially reversible by *Pkd1* reactivation(43), the persistent effect of *Pkd1* deletion at each PKD stage may be critical for PKD progression, underscoring the importance of analysis at time-course analysis following *Pkd1* deletion. Comparing and contrasting with the mouse PKD atlas to human ADPKD dataset (13) shed light on the conserved cellular heterogeneity (Fig. 2-4) in PKD as well as shared molecular mechanism including transcriptional regulation of a cyst lining cell marker; GPRC5A (Fig. 5).

Our single cell analysis indicated that the differentially expressed genes as well as accessible regions in *Pkd1*-deficient tubular epithelia are highly cell-type-specific (Fig. S6), although the resultant phenotype; cyst formation is shared among tubular compartments. This observation suggests that the response to *Pkd1* deletion as well as the molecular mechanism of cyst formation may be cell-type-specific. Furthermore, the effect of *Pkd1* deletion in one cell type may affect the responses to *Pkd1* deletion in other cell types through modifying the microenvironment, potentially further complicating the molecular response to *Pkd1* deficiency in vivo. In addition, we found that the cell-type-specific response to *Pkd1* is dynamically changed along PKD progression (Fig. 2E, F, Fig. 3G). The medullary principal cell subpopulation initially showed molecular signature consistent with DNA damage response and alteration of metabolic pathways following *Pkd1* deletion, and subsequently inflammatory responses are prominent at a later time point (Fig. 2E). Our findings emphasize the importance of single cell analysis with a time course for dissection of PKD mechanism.

In this study, we described that heterogeneous FR-PTCs with distinct molecular signatures are components of proximal tubular cysts. Furthermore, FR-PTCs in PKD show unique gene expression signature compared to those observed after ischemia-reperfusion injury (Fig. S15) (10). Pseudotemporal analysis indicated FR-PTC subtypes separately arise from a transitional cell state, and their chromatin accessibilities as well as transcription factor activities are unique (Fig. 4). FR-PTC1 showed RELA and AP-1 family activation and subsequent inflammatory molecular signature, while FR-PTC2 is characterized by TEAD family activation probably through Hippo pathway deregulation (Fig. 4). However, the differential roles of these heterogeneous FR-PTCs in PKD progression have remained elusive in this study. Future genetically engineered approaches and lineage-tracing in a mouse PKD model may reveal their contributions to PKD progression.

Our previous single cell analysis on human ADPKD identified GPRC5A as a cyst lining cell marker with epigenetic deregulation in ADPKD(13). Interestingly, GPRC5A is a cyst marker also in mouse kidneys regardless of their proximal or distal origin (Fig. 5). The 5’ distal enhancer activating GPRC5A expression in human ADPKD was conserved in mice (Fig. 5). The 5’ distal enhancer is characterized by multiple TEAD binding motifs both in mouse (Fig. S16) and human kidneys(13). Indeed, Hippo pathway deregulation by LATS inhibition up-regulated GPRC5A expression in human primary PTC (Fig.5). This finding suggests that up-regulation of GPRC5A expression is involved in Hippo pathway deregulation in cystic epithelia. The regulation of GPRC5A expression by Hippo pathway is in line with the previous study implicating Hippo pathway in PKD progression(36). Our finding suggests that the disturbance of the gene regulatory networks driving GPRC5A expression may be shared with multiple cell types in mouse and human PKD despite the general cell-type-specificity of the responses to *Pkd1* deletion (Fig. S6). However, the function of GPRC5A as well as physiological ligand in vivo remains elusive. Future study with a PKD mouse model with *Gprc5a* deletion or pharmacologic inhibition may reveal the role of GPRC5A in cyst growth in PKD.

In summary, we performed multiomic single nucleus analysis for a mouse PKD to characterize temporally dynamic, cell-type-specific deregulation of gene expressions and chromatin accessibilities. By comparing our mouse PKD datasets with human ADPKD datasets, our study sheds light on the conservation of cellular heterogeneity in PKD. Our findings collectively indicated that the intersectional analysis for our mouse and human PKD dataset is useful approach to describe PKD mechanism and identify potential therapeutic targets. This study is limited by the lack of experimental analysis on the roles of heterogeneous principal cells and FR-PTC subtypes in PKD progression. Furthermore, the mechanistic role of GPRC5A in PKD progression needs to be addressed. Despite these limitations, our single nucleus multimodal analysis of mouse PKD as well as our analytic approach provide a foundation on which to promote future efforts to understand cell-specific mechanism of cystic tubular changes and develop better therapeutic approaches for PKD.

## Materials and Methods

### PKD mouse model

All animal experimental procedures were conducted following the University of Maryland or Washington University Animal Care and Use Committee (IACUC) guidelines and procedures. The *Pkd1* conditional allele (*Pkd1*^*fl/fl*^) used in these studies has been previously described(15). The *Pkd1*^*fl/fl*^; *Pax8*^*rtTA*^; *TetO*^*Cre*^ mouse model is an orthologous mouse model of ADPKD that utilizes the murine *Pax8* promoter to drive expression of the reverse tetracycline-dependent transactivator (rtTA) to all renal tubular compartments (14). To induce *Pkd1* deletion, experimental mice were treated with a single daily intraperitoneal injection of doxycycline hyclate (Sigma-Aldrich, D9891) 50 mg/Kg body weight, diluted in sterile distilled water. Specific time points for induction are indicated P27, P28, P29, P44 and P56. Littermate controls lacked either the *Pax8*^*rtTA*^ or the *TetO*^*Cre*^ allele but were treated with doxycycline in the same manner as experimental mice. All mice were maintained on an inbred C57BL/6 background. Males were included in this study. Kidneys were harvested at P66, P100 and P130. The right kidneys were decapsulated and snap-frozen in liquid nitrogen for single-nucleus multiomics analysis. The kidneys for *Pkd1*^*fl/fl*^; *Six2*^*Cre*^ mouse model were analyzed at P7. The *Pkd1*p.R3269C knock-in mice (equivalent to human *Pkd1*p.R3277C mutation [*Pkd1*^RC/RC^])(40, 44) were analyzed at 11 months of age.

### Mouse genotyping

Genomic DNA was isolated from mouse tails or toes and genotyped using the primers on the Table S4. The genotyping PCR was performed with the following cycles; 56°C, 63°C, and 59°C annealing temperature and 35 PCR cycles for *Pkd1* ^F/F^ and *Pax8*^*rtTA*^ and *TetO*^*Cre*^ transgenes respectively. The PCR products were resolved on agarose gels, yielding a product of ∼250 base pairs (bp), ∼595 bp, and ∼235 bp for *Pkd1*^*fl/fl*^ and *Pax8*^*rtTA*^ and *TetO*^*Cre*^ transgenes, respectively.

### Nuclear dissociation for library preparation

For single nucleus sequencing, nuclei were isolated with Nuclei EZ Lysis buffer (NUC-101; Sigma-Aldrich) supplemented with protease inhibitor (5892791001; Roche) and RNase inhibitor (Protector RNase Inhibitor). Samples were cut into < 1 mm pieces, homogenized using a Dounce homogenizer (885302–0002; Kimble Chase) in 2 ml of ice-cold lysis buffer, and incubated on ice for 5 min with an additional 1 ml of lysis buffer. The homogenate was filtered through a 40-μm cell strainer (43–50040–51; pluriSelect) and centrifuged at 500 g for 5 min at 4°C. The pellet was resuspended, washed with 3 ml of buffer, and incubated on ice for 5 min. Following centrifugation, the pellet was resuspended in Nuclei Buffer (10× Genomics, PN-2000153), filtered through a 5-μm cell strainer (43-50005-03, pluriSelect), and counted. All these processes were performed in a cold room at 4°C. Subsequently, nuclei suspension was diluted to target 10,000 nuclei per lane and loaded into a thermal cycler to allow transposition reaction.

### Single nucleus multiomics library sequencing and following bioinformatics workflow

Single-nucleus multiomics libraries were generated using Chromium Next GEM Single Cell Multiome ATAC+ Gene Expression Reagent Kit (10X Genomics) following nuclear dissociation. Libraries were sequenced on an Illumina Novaseq instrument and counted with CellRanger ARC v2.0.2 (10X Genomics) using refdata-cellranger-arc-mm10-2020-A-2.0.0, which is mm10 reference genome 10X Genomics provided. The read configuration was 2x150bp paired-end. The output of CellRanger ARC was processed through Seurat v4.0.2(16). Each of datasets was processed to subset high-quality nuclei (nCount_RNA > 1000 & nCount_RNA < 15000 & nFeature_RNA > 500 & nFeature_RNA < 4000 and those with %Mitochondrial genes < 0.25 for snRNA-seq data; nCount_ATAC < 40000 & nCount_ATAC > 1000 & nucleosome_signal < 2 and those with TSS enrichment score > 1 for snATAC-seq data). Heterotypic doublets were identified with DoubletFinder v2.0.3 (45) as instructed, and the estimated doublets were removed after merging datasets. The datasets were aggregated in Seurat using the merge function. snATAC-seq peaks on the aggregated data were called with MACS2 (v2.2.7.1) using CallPeaks function in Signac with default parameters and a new Seurat object was generate from these peaks using FeatureMatrix and CreateChromatinAssay functions, with annotation generated by GetGRangesFromEnsDb(ensdb = EnsDb.Mmusculus.v79). Following normalization with SCTransform with regressing out the %mitochondrial reads and cellular cell-cycle status (cell-cycle scores calculated by CellCycleScoring function), batch effects were corrected with Harmony (17) using the RunHarmony function in Seurat on the SCT assay. To process the snATAC-seq data term-frequency inverse-document-frequency (TFIDF) was computed, then dimensional reduction was performed on the TFIDF matrix with singular value decomposition. Harmony was used for batch effect correction for snATAC-seq clustering. Finally, a weighted nearest neighbor (WNN) graph was constructed with a weighted combination of snRNA-seq and snATAC-seq data with the FindMultiModalNeighbors function. Clustering was performed by small local moving (SLM) with the FindCluster function (resolution of 0.6). Dimensional reduction was performed with the RunUMAP function and individual clusters were annotated based on the expression of lineage-specific markers.

The aggregated dataset was composed of 125,434 nuclei (n=18 mice) with 66,345 nuclei for control (n=9) and 59,089 for PKD (n=9). There was a mean of 8293 ± 1031 nuclei per sample for control and of 7386 ± 710 nuclei per sample for PKD. There was a mean of 3875 UMIs, 1958 genes and 7190 peaks detected per nucleus for control. There was a mean of 3966 UMIs, 2023 genes and 7587 peaks detected per nucleus for control (Fig. S2, S3).

Differential expression between cell types was assessed with the FindMarkers function for transcripts detected in at least 10% of cells with a log-fold-change threshold of 0.25 (11-13). Differential accessible regions (DARs) were assessed with the Seurat FindMarkers function for peaks detected in at least 10% of cells with a likelihood ratio test and a log-fold-change threshold of 0.2 and test.use = ‘LR’, latent.vars = “nCount_peaks”. Bonferroni-adjusted p-values were used to determine significance at an FDR < 0.05.

### Sub-clustering analysis

The distal nephron clusters (CNT, PC1, PC2 and URO; Fig. 2) or FRPTC/PEC (Fig. 3) were subset from the whole dataset and performed sub-clustering. The snRNA-seq data were normalization with SCTransform with regressing out the %mitochondrial reads and cellular cell-cycle status (cell-cycle scores calculated by CellCycleScoring function), For the snATAC-seq data, term-frequency inverse-document-frequency (TFIDF) was computed, then dimensional reduction was performed on the TFIDF matrix with singular value decomposition. Finally, a weighted nearest neighbor (WNN) graph was constructed by identifying the nearest neighbors. Clustering was performed by small local moving (SLM) algorithm with FindCluster function. Dimensional reduction was performed with the RunUMAP function and individual clusters were annotated based on the expression of lineage-specific markers. For distal nephron analysis, the remaining low QC clusters with other cell type markers were further removed after sub-clustering.

### Estimation of transcription factor activity from snATAC-seq data

Transcription factor binding motif enrichment was assessed on the aggregated dataset with chromVAR v1.10.0(19). The positional weight matrix was obtained from the JASPAR2022 database (collection = “CORE”, tax_group = ‘vertebrates’, all_versions = F) (46). Cell-type-specific chromVAR activities were calculated using the RunChromVAR function (Signac v1.4.0). The chromVAR motif enrichment score for each transcription factor on the whole dataset was visualized with FeaturePlot function with max.cutoff = q99 and min.cutoff = q1 or VlnPlot function. Differentially enriched motifs were assessed with the FindMarkers function for with a log-fold-change threshold of 0.1 and mean.fxn = rowMeans, fc.name = “avg_diff”.

### Visualization of single nucleus dataset features

Gene expressions (snRNA-seq, assay SCT) were visualized with FeaturePlot (UMAP), VlnPlot (violin plot) or DotPlot (dot plot) function on Seurat. The fragment coverage around differentially accessible regions in each cell type was visualized by CoveragePlot function (Signac).

### Construction of pseudotemporal trajectories

Pseudotemporal trajectories for the snRNA-seq were constructed on Monocle 3 according to instructions provided on GitHub (https://cole-trapnell-lab.github.io/monocle3)(47–49). Briefly, raw count matrix subset for FRPTC1/2/3 and transitional PTC in FRPTC_PEC subclutering (Fig. 3A) was used to generate cell data set object (CDS) with “new_cell_data_set” function. Next, the CDS was preprocessed (num_dim = 30, method = “PCA”), aligned to remove batch effect among samples(50) and reduced onto a lower dimensional space with the “reduce_dimension” function (preprocess_method = “Aligned”). The cells were then clustered (“cluster_cells”) and ordered with the “learn_graph” function. We used the “order_cell” function and indicated the transitional PTC as “start point” of the trajectories.

### Generation of cis-coaccessibility network

Cis-coaccessibility networks (CCAN) were predicted using Cicero v1.3.5 according to instructions provided on GitHub (https://cole-trapnell-lab.github.io/cicero-release/docs_m3/)(51). Briefly, the human ADPKD kidneys(13) or mouse FRPTC subtypes in PKD kidneys (Fig. 3A) were extracted and converted to cell dataset (CDS) objects using the make_atac_cds function. The CDS object was processed by the detect_genes() and estimate_size_factors() functions, followed by dimensional reduction. The Cicero connections were obtained using the run_cicero function with default parameters. CCAN was visualized with plot_connections function (coaccess_cutoff = .1). The CRE for human GPRC5A in hg38 (chr12:12871973-12873059) was lifted to mm10 (chr6:135040191-135040827) with Lift Genome Annotations (Minimum ratio of based that must remap = 0.5) in UCSC Genome Browser (https://genome.ucsc.edu/cgi-bin/hgLiftOver) (52).

### Cell culture

Human primary proximal tubular cells (human primary PTC, Lonza; CC-2553) were cultured with renal epithelial cell growth medium kit (Lonza; CC-3190) in a humidified 5% CO2 atmosphere at 37°C. Experiments were performed on early passages. Cells were plated at a density of 1×10^5^ cells per well in a 6-well plate, incubated overnight, and transfected with 50 nM siRNA for *LATS1* and *LATS2* (ON-TARGETplus SMARTpool siRNA [Horizon Discovery]) or negative control siRNA (ON-TARGETplus Non-targeting Control Pool [Horizon Discovery]) using Lipofectamine RNAiMAX (Life Technologies) following the manufacturer’s protocol. Cells were harvested at 72 h after transfection for RNA isolation.

### Quantitative PCR

Total RNA was extracted from primary human PTC with the RNeasy Mini kit (Qiagen) following manufacturer’s instructions. The extracted RNA was reverse transcribed using the High-Capacity cDNA Reverse Transcription Kit (Life Technologies). Quantitative PCR (qPCR) was performed using iTaq Universal SYBR Green Supermix (Bio-Rad). Data were normalized by the abundance of *GAPDH* mRNA. The following primers were used: *GAPDH*: Fw 5′-GACAGTCAGCCGCATCTTCT -3′; Rv 5`-GCGCCCAATACGACCAAATC -3′; *GPRC5A*: Fw 5′-ATGGCTACAACAGTCCCTGAT -3′; Rv 5`-CCACCGTTTCTAGGACGATGC -3′.

### Reanalysis of ChIP-seq data

The ChIP-seq dataset for TEAD1 binding in MCF7 or T47D cells were retrieved from GSE107013 (42). The CRE for human GPRC5A in hg38 (chr12:12871973-12873059) was lifted to hg19 (chr12:13024907-13025993) with Lift Genome Annotations (Minimum ratio of based that must remap = 0.95) in UCSC Genome Browser (https://genome.ucsc.edu/cgi-bin/hgLiftOver) (52). The peaks on the lifted CRE was visualized with the downloaded BigWig files and IGV.

### Gene set enrichment analysis and gene ontology analysis

Single nucleus gene set enrichment analysis was performed with the VISION v2.1.0 R package according to instructions provided on GitHub (https://github.com/YosefLab/VISION)(53), using Hallmark gene sets as well as Hippo pathway gene set (REACTOME-SIGNALING-BY-HIPPO) were obtained from the Molecular Signatures Database v7.4 distributed at the GSEA Web site. The heatmaps were generated with pheatmap v1.0.12 from gene set enrichment scores averaged in each subtype.

### Immunofluorescence studies

Mice were euthanized, and left kidneys were fixed with 4% formaldehyde (Pierce, #28908) overnight and embedded in paraffin and sectioned. Paraffin-embedded tissue sections were deparaffinized by immersing glass slides with sections into xylene and ethanol. The slides were placed in antigen retrieval solution (Vector H-330) and incubated in a pressure cooker. The sections were blocked in blocking buffer [1% BSA (Roche; 03116956001), 0.1% Triton X-100 (Sigma; T8787) in PBS] for 1 h, and the following primary antibodies or glycoproteins were added (1:200); LTL (Vector Labs; B-1325 or FL-1321), DBA (Vector Labs; B-1035 or FL-1031), sheep Anti-UMOD (Bio-Rad; 8595-0054), rabbit Anti-Calbindin1 (Abcam; ab108404), goat Anti-AQP2 (Novus Biologicals; NBP1-70378), rabbit Anti-TMSB4X (Immunodiagnostik, A9520.2), goat Anti-VCAM1 (R&D Systems; AF643-SP), rabbit Anti-VCAM1 (Abcam; ab134047), rabbit Anti-SPP1(Abcam, ab218237)]), mouse Anti-CFTR (Cystic Fibrosis Foundation, 596), rabbit Anti-GPRC5A (Novus Biologicals; NBP1-89743) and incubated overnight at 4°C. When CFTR was stained, The next day, slides were wash with PBS, and secondary antibodies in blocking buffer were added (1:200) [donkey Anti-Goat (Invitrogen; A11057 or A32814), Donkey Anti-Mouse (Invitrogen; A10037), donkey Anti-Rabbit (Invitrogen; A10042 or Jackson ImmunoResearch; 711-545-152), Anti-Sheep antibody (Jackson ImmunoResearch; 713-165-147), conjugated Streptavidin (Invitrogen; S21374) and incubated at room temperature for 1 h. Sections were stained with DAPI (4 ',6' -diamidino-2-phenylindole) and mounted in Prolong Gold (Life Technologies). Imaging was performed on a Nikon Eclipse Ti Confocal at 20X objective and processed using Nikon Elements-AR. Imaging conditions (exposure time, laser intensity) and processing (background subtraction, color balance) were optimized for each antibody.

## Statistical analysis

No statistical methods were used to predetermine sample size for single nucleus analysis. Experiments were not randomized and investigators were not blinded to allocation during library preparation, experiments or analysis. Bonferroni adjusted *P* values were used to determine significance for differential accessibility. The quantitative PCR data (Fig. 5) are presented as mean±s.d. and were compared between groups with two-sided Student’s *t*-test. The kidney to body weight ratios (Fig. 1B) were presented as mean±s.d. and one-way ANOVA with post hoc Turkey test. A *P* value of <0.05 was considered statistically significant.

## Acknowledgments

These experiments were funded by NIDDK UC2DK126024, 2R01DK103740 and 1U54DK137332 (B.D.H.). Additional support was from American Society of Nephrology Carl W. Gottschalk Research Scholar Award and the Pilot and Feasibility Grant of Polycystic Kidney Disease Research Resource Consortium (U24DK126110) (Y.M.); and the Department of Defense W81XWH-20-1-0198 (M.R.M.). The CFTR antibody was provided by Martina Gentzsch, Ph.D., University of North Carolina, Chapel Hill, and Cystic Fibrosis Foundation.

## Author Contributions

Y.M. performed experiments with contributions from Y.Y., M.C.P. Y.M., H.W., and N.L. performed bioinformatic analysis. Y.M., Y.Y., H.W., M.C.P., N.L., O.M.W, P.G., T.J.W., and B.D.H. analyzed data. Mouse PKD samples were collected by P.G., T.C., M.R.M. Y.M. and B.D.H. designed experiments and wrote the manuscript. All authors read and approved the final manuscript.

