## Supplemental Materials for "Multi-omics profiling of mouse polycystic kidney disease progression at a single cell resolution"

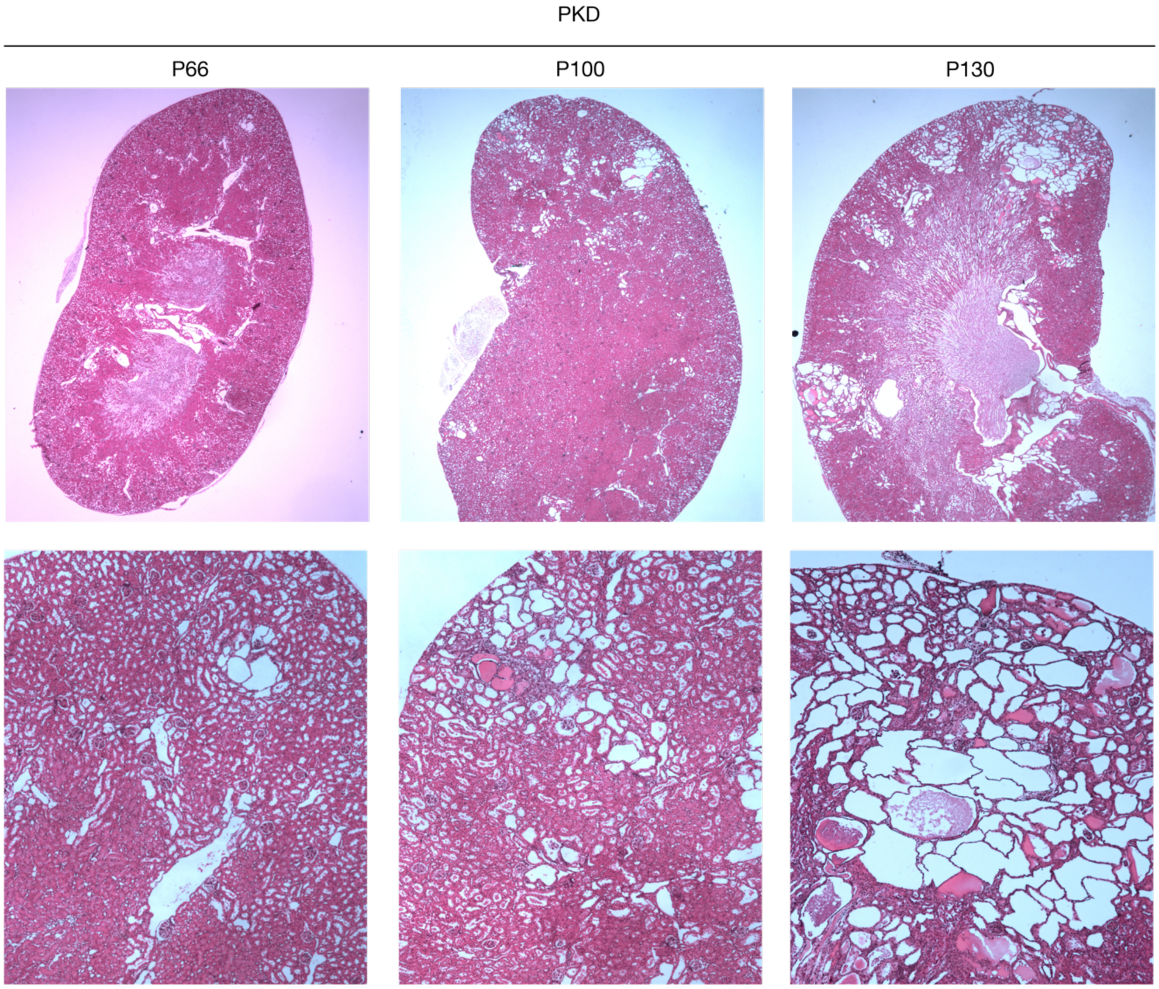


**Fig. S1. Appearance of kidneys in PKD model mice**: Representative H.E. staining images of the kidneys from *Pkd1*$^{fl/fl}$*; Pax8*$^{rtTA}$*; TetO*$^{Cre}$ mice at P66, P100 or P130.

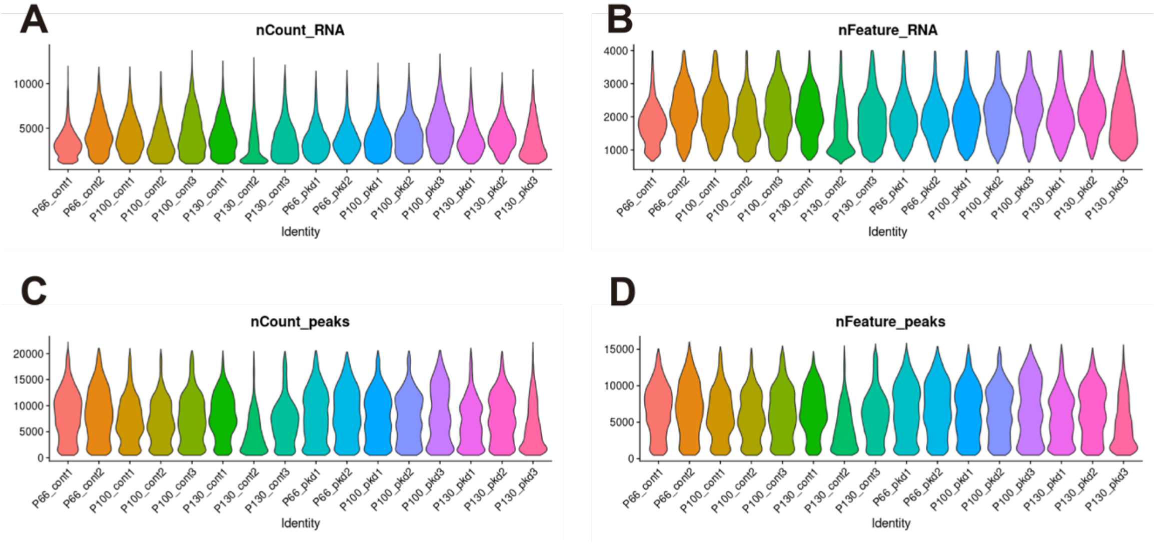


**Fig. S2. QC metrics of single nucleus multiomics data for each sample**: Violin plot showing; (A) number of UMI, (B) number of genes detected per nucleus for individual samples in snRNA-seq. Violin plot showing; (C) number of fragments, (D) number of peaks per nucleus for individual samples in snATAC-seq.

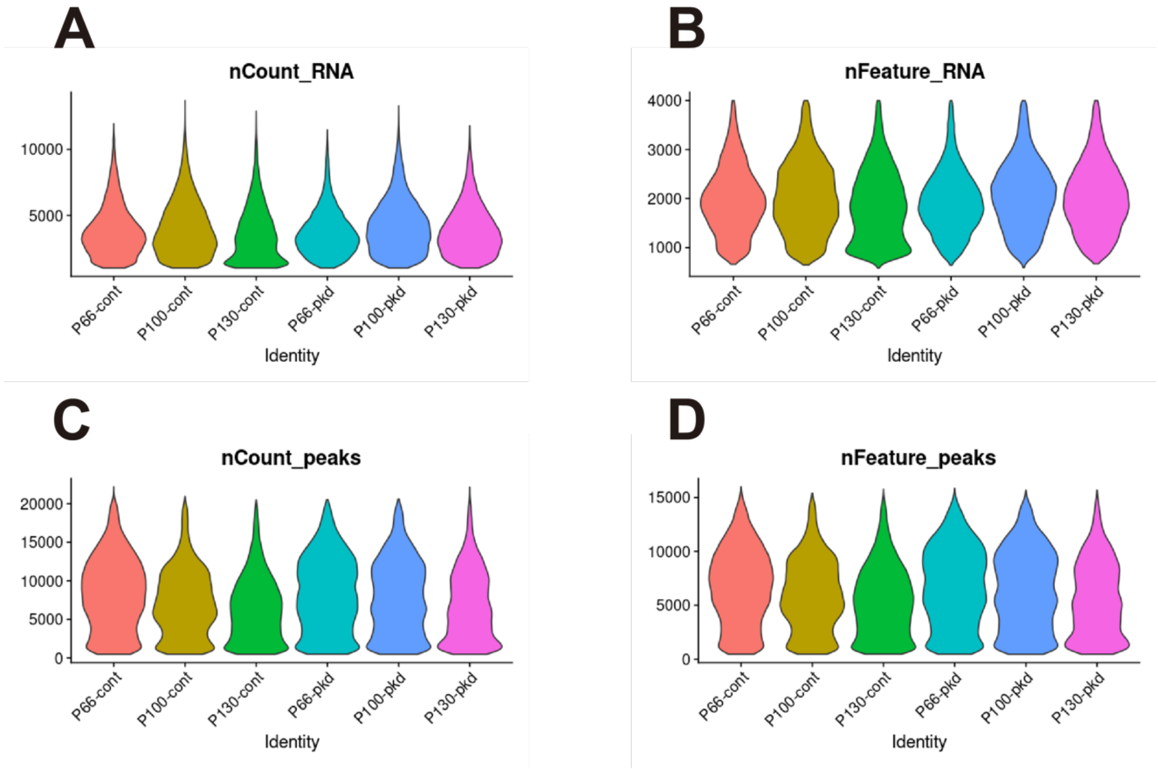


**Fig. S3. QC metrics of single nucleus multiomics data for each condition**: Violin plot showing; (A) number of UMI, (B) number of genes detected per nucleus for each condition in snRNA-seq. Violin plot showing; (C) number of fragments, (D) number of peaks per nucleus for each condition in snATAC-seq.

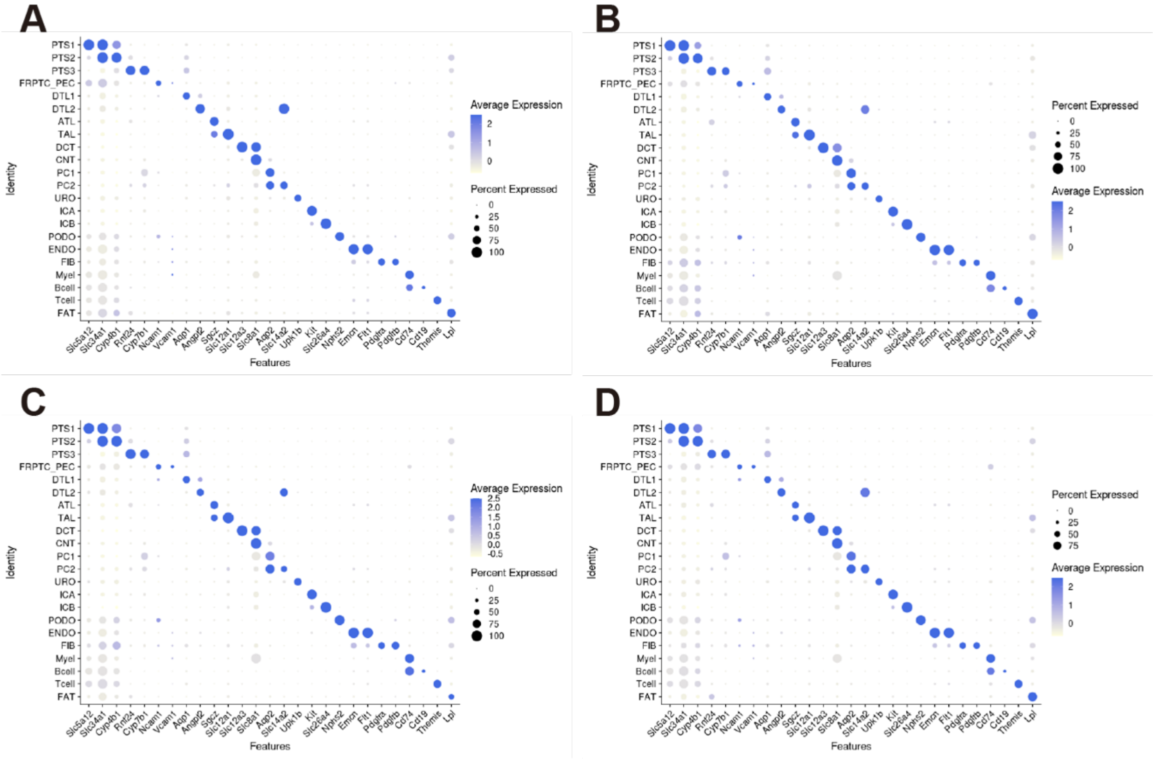


**Fig. S4. Lineage marker gene expressions in each PKD stage**: Dot plot showing gene expression patterns of cluster-enriched markers for the integrated dataset of control (A), PKD at P66 (B), P100 (C) or P130 (D). The diameter of the dot corresponds to the proportion of cells expressing the indicated gene and the density of the dot corresponds to average expression relative to all cell types.

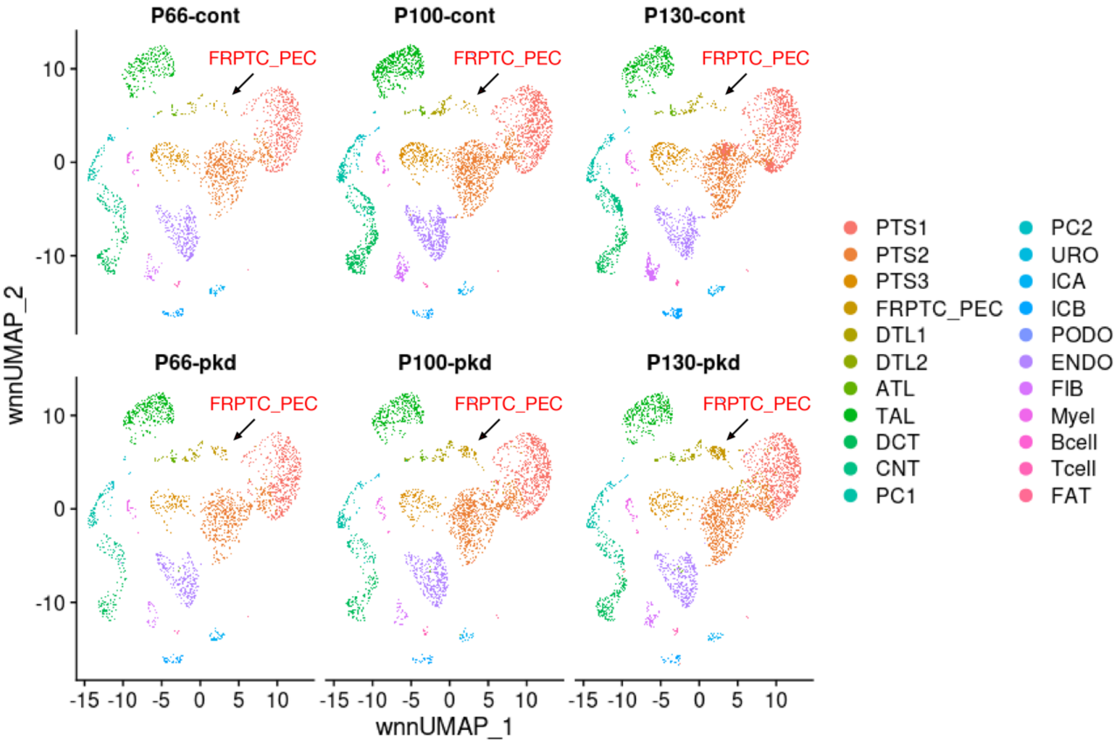


**Fig. S5. UMAP plot of the integrated dataset separately shown for each condition**: UMAP plot of the whole dataset split by conditions. The arrows indicate the FRPTC_PEC cluster.

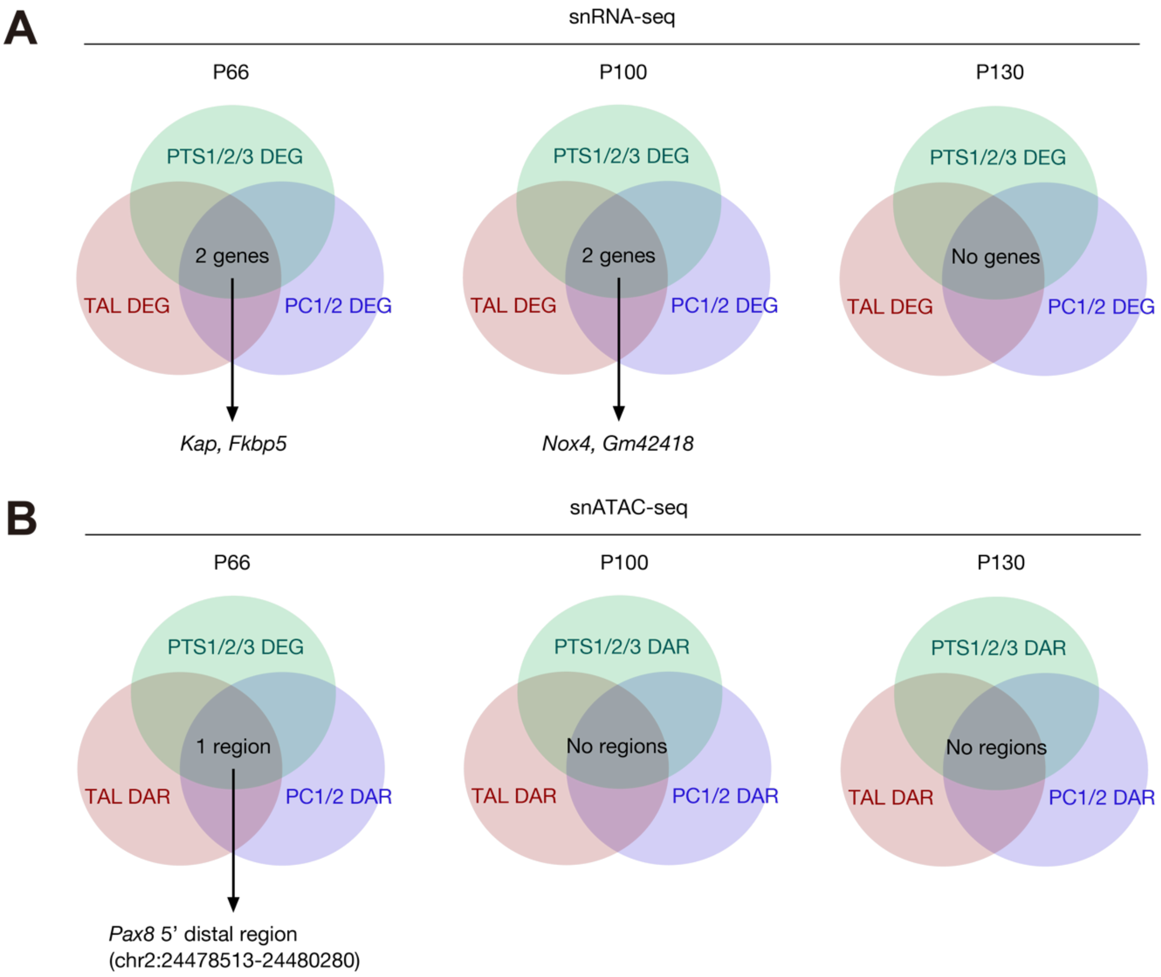


**Fig. S6. Shared differentially expressed genes or accessible regions among major tubular cell types**: (A) Shared PKD DEGs among PTS1/2/3, TAL and PC1/2 at P66, P100 or P130. (B) Shared PKD DAR among PTS1/2/3, TAL and PC1/2 at P66, P100 or P130.

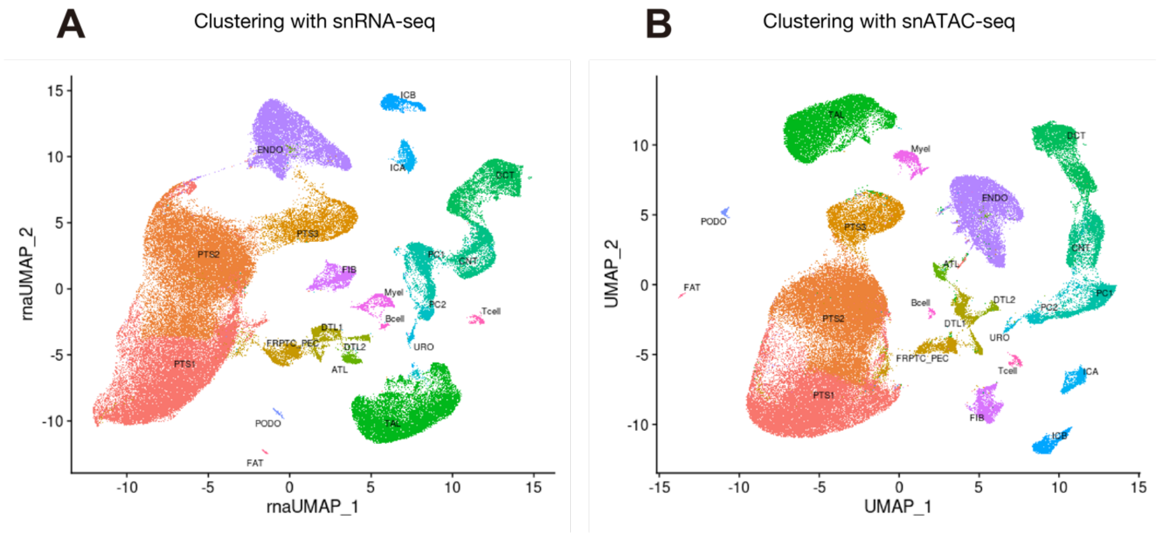


**Fig. S7. UMAP plot of the integrated dataset clustered based on each modality**: UMAP plot of the whole dataset clustered by snRNA-seq (A) or snATAC-seq (B).

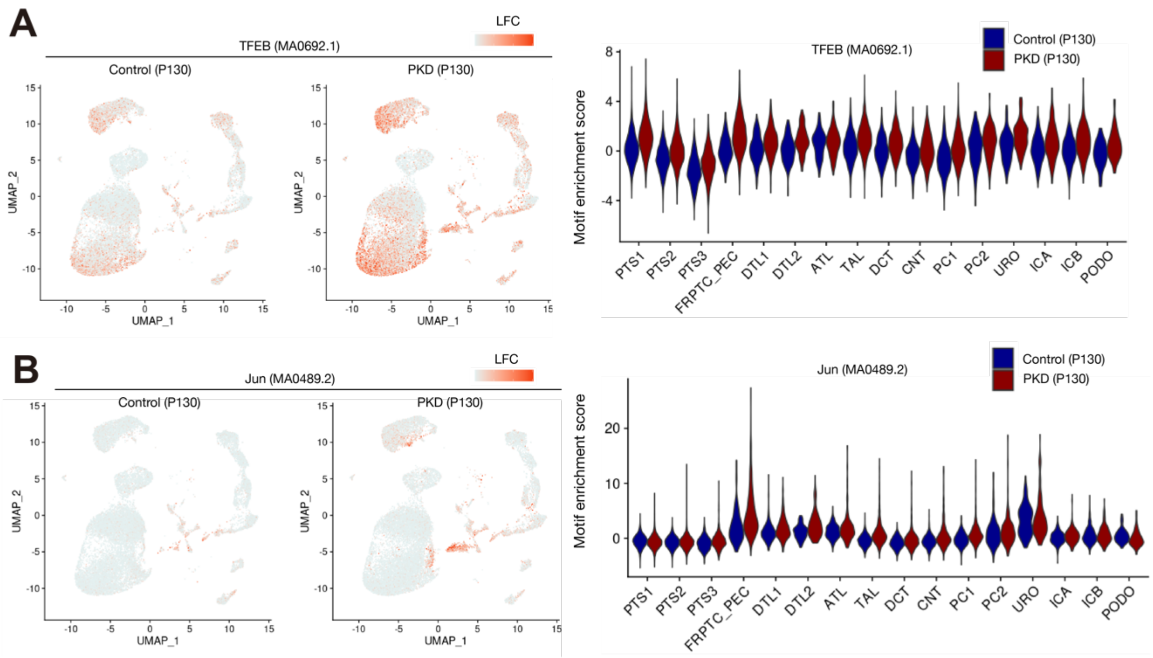


**Fig. S8. TFEB and AP-1 motif enrichment in mouse PKD cell types**: UMAP displaying enrichment of transcription factor binding motifs in control or PKD kidneys (left) or violin plot showing the motif enrichment scores in each cell type (right) for (A) TFEB (MA0692.1) or (B) Jun (MA0489.2). The color scale of UMAP plot represents a normalized log-fold-change.

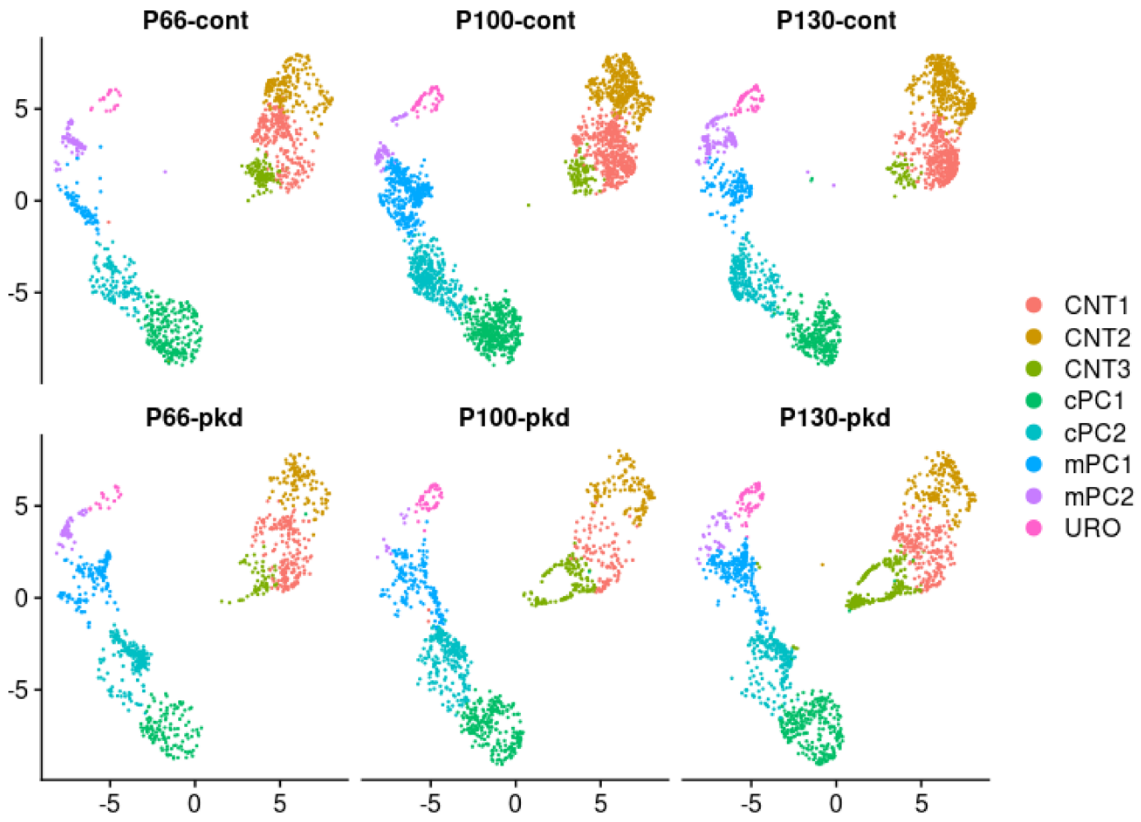


**Fig. S9. UMAP plot of the distal nephron subclustering separately shown for each condition**: UMAP plot of the distal nephron subtypes split by conditions. All the subtypes are detected in each condition.

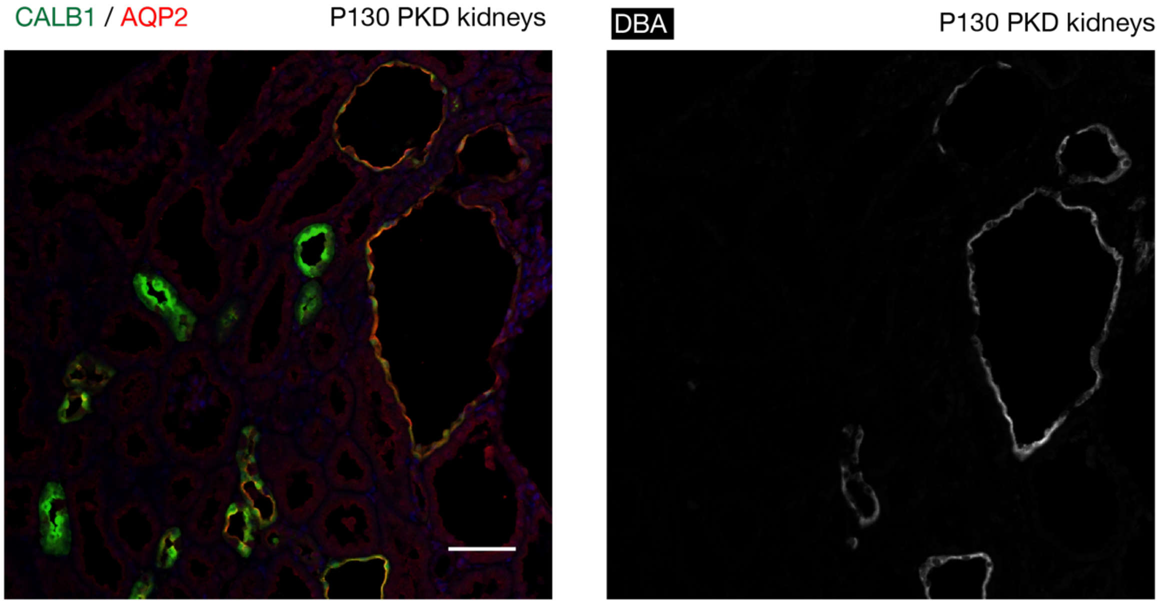


**Fig. S10. AQP2+ collecting duct lineage constitutes DBA+ cyst epithelia**: Representative immunofluorescence images of CALB1 (green), AQP2 (red) and DBA (white) in the PKD kidneys at P130. Scale bar indicates 50 μm.

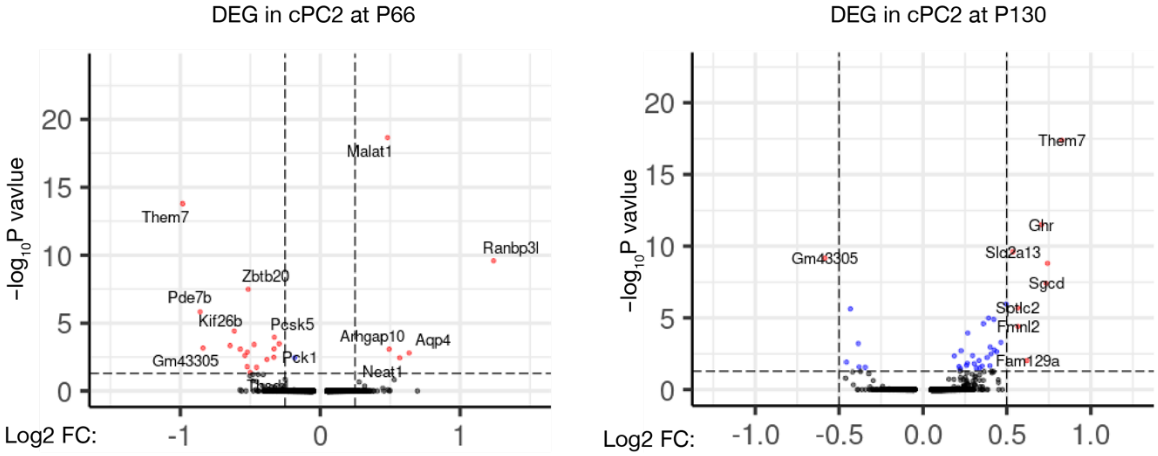


**Fig. S11. Differentially expressed genes in cPC2 subtype**: Volcano plot showing differentially expressed genes in cPC2 of PKD mice compared with that of control at P66 (left) and P130 (right).

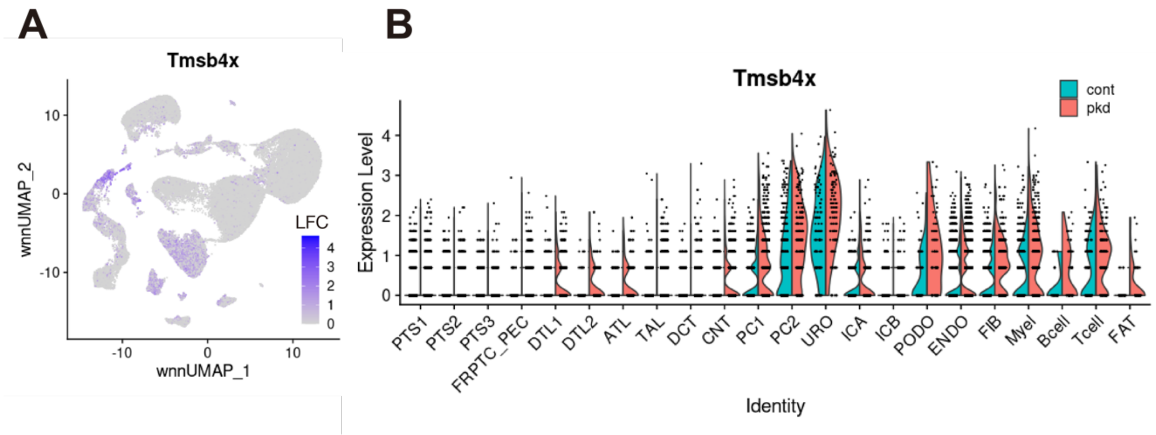


**Fig. S12. *Tmsb4* expression in the P130 control and PKD dataset**: Feature plot (A) or Violin plot (B) showing the expression levels of *Tmsb4x* for control or PKD at P130 in snRNA-seq.

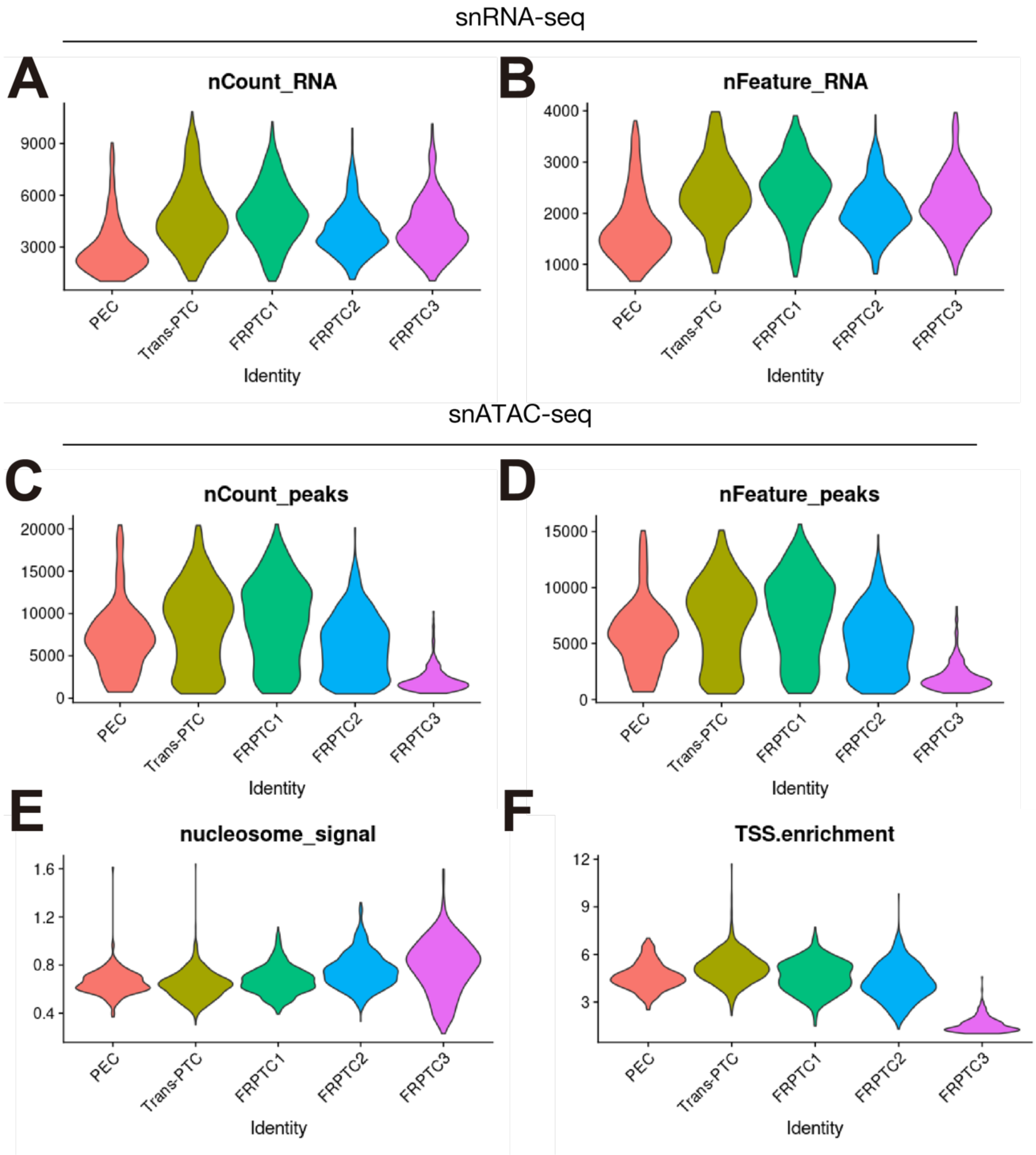


**Fig. S13. QC metrics among subtypes in PEC/FRPTC cluster**: Violin plot showing; (A) number of UMI, (B) number of genes detected per nucleus for each subtype in snRNA-seq. Violin plot showing; (C) number of fragments, (D) number of peaks, (E) nucleosome signal or (F) TSS enrichment score per nucleus for each subtype in snATAC-seq.

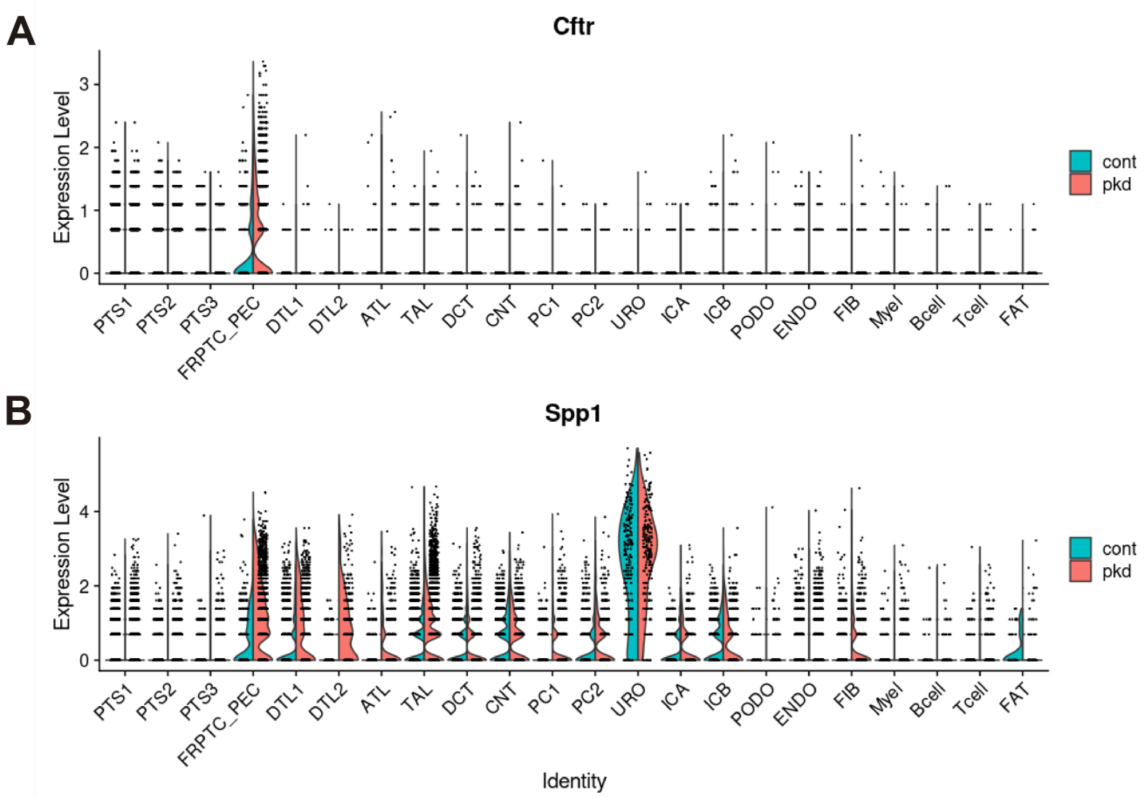


**Fig. S14. *Cftr* and *Spp1* expression in the whole dataset**: Violin plot showing the expression levels of (A) *Cftr* or (B) *Spp1* for control or PKD in snRNA-seq.

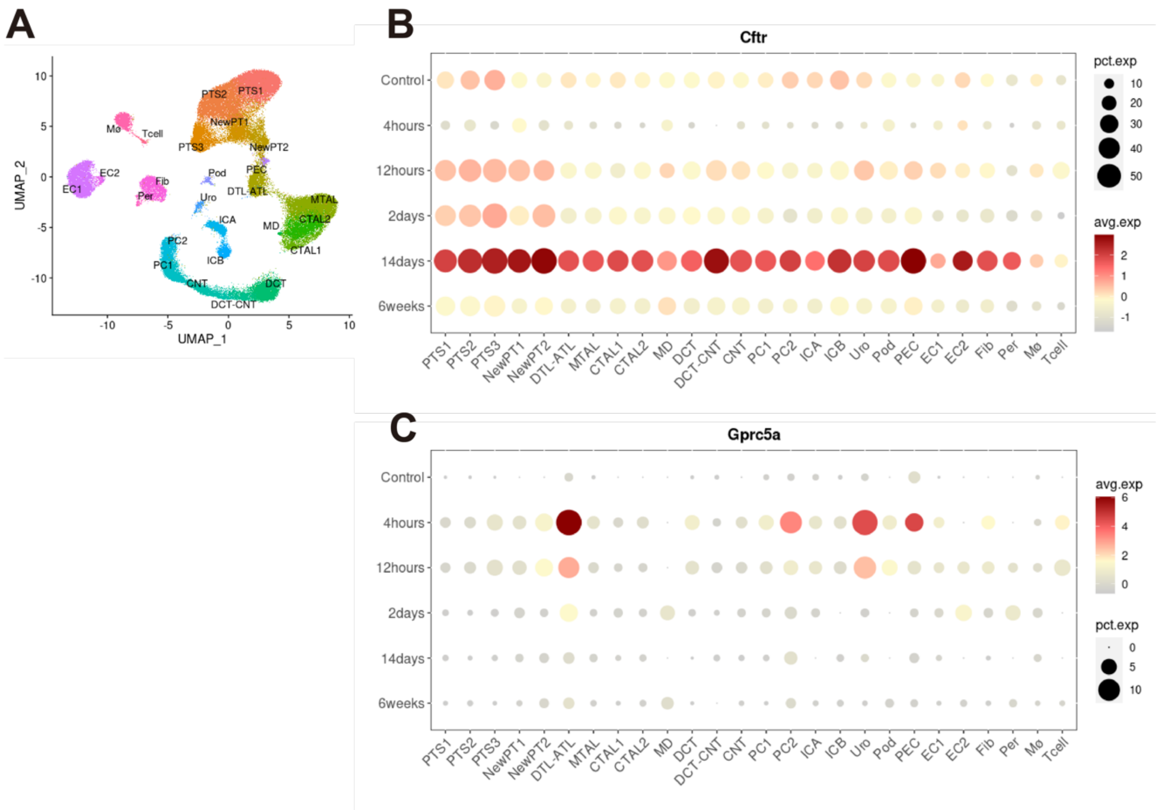


**Fig. S15. *Cftr* and *Gprc5a* expression in the mouse kidneys with ischemia-reperfusion injury**: Reanalysis of published snRNA-seq for the mouse kidneys with ischemia-reperfusion injury (GSE139107). UMAP and annotations were shown in (A). Dot plot showing the expression of (B) *Cftr* or (C) *Gprc5a* in each cell type at each time point following ischemia-reperfusion injury. The diameter of the dot corresponds to the proportion of cells expressing the indicated gene and the density of the dot corresponds to average expression relative to all cell types.

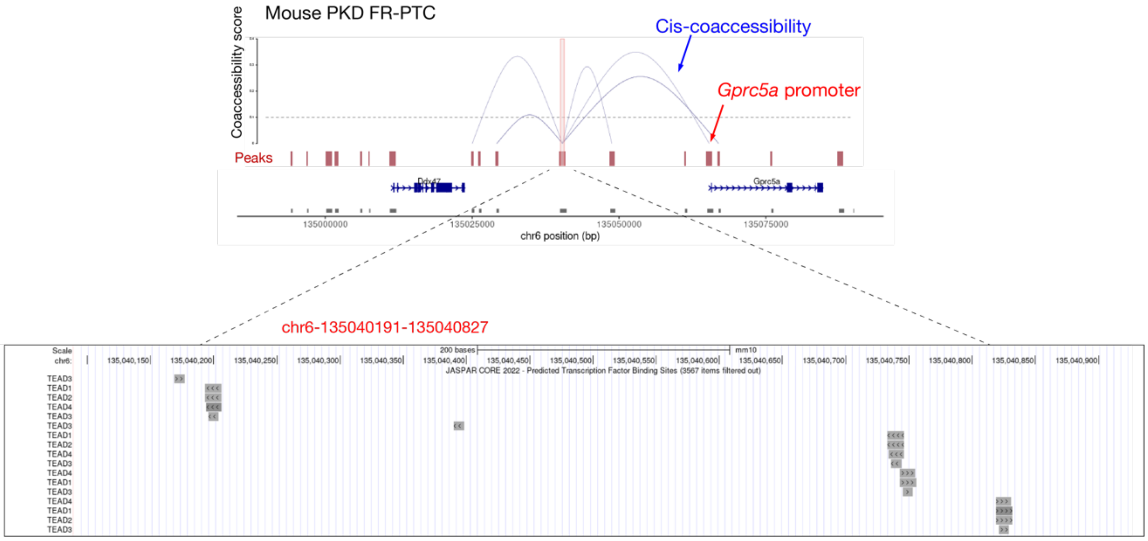


**Fig. S16. TEAD family binding motifs in the cis-regulatory region of *Gprc5a***: The 5' distal cis-regulatory region (CRE) of *Gprc5a* gene in PKD mouse model (upper panel, redundant with Fig. 5G) has several binding motifs for TEAD family transcription factors (TEAD1-4), based on JASPAR CORE vertebrates collection (2022) shown on the UCSC genome browser (Enrichment score > 300, lower panel).

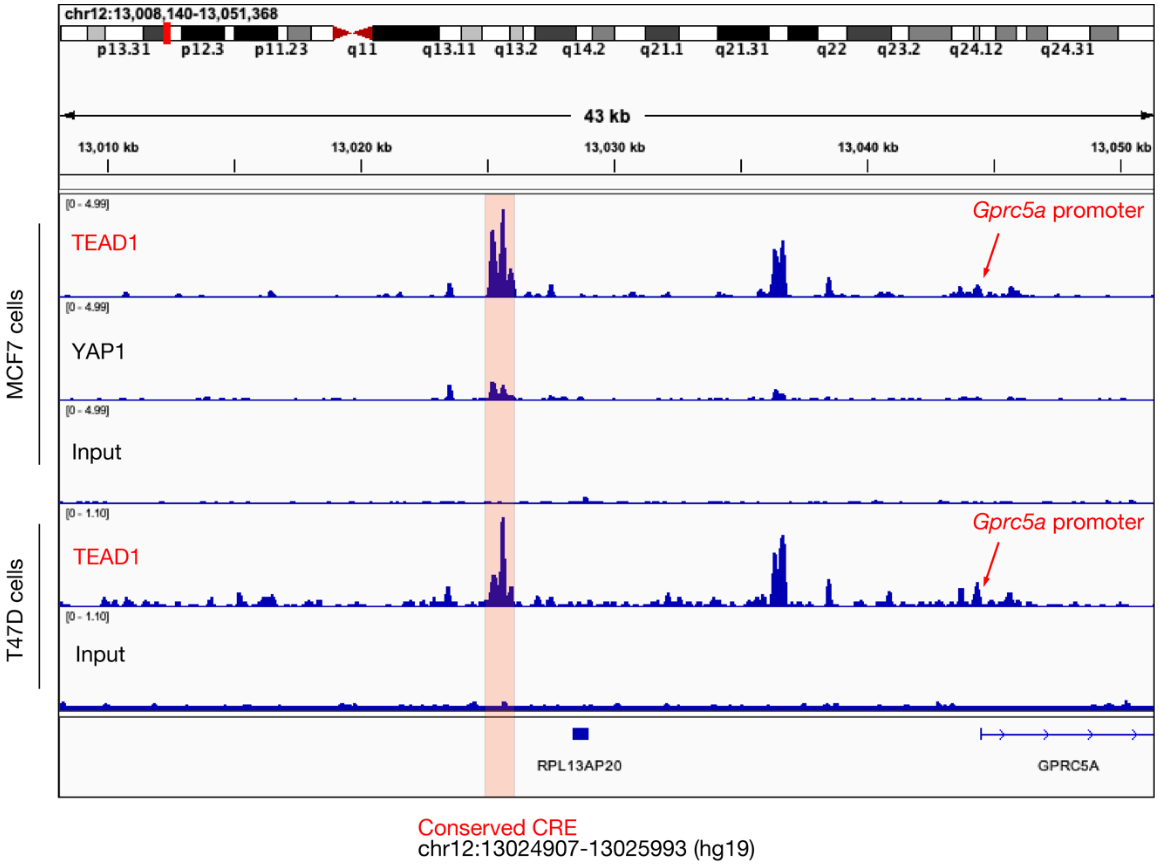


**Fig. S17. TEAD1 binds to the cis-regulatory region of GPRC5A.** ChIP-seq data visualization of TEAD1 and YAP1 binding sites around the 5' distal regions of *GPRC5A* gene in human genome (hg19). The conserved cis-regulatory region (CRE) of *GPRC5A* gene was shown to bind with TEAD1 in two human breast cancer cell lines.

| Library ID | Genotype | Disease | Time | KW | BW | KW/BW |
|---|---|---|---|---|---|---|
| P066_cont1 | Pkd1^neo;Pax8-;TET-OCre+ | P066_control | p66 | 0.1255 | 19.6 | 0.00640306 |
| P066_cont2 | Pkd1^neo;Pax8-;TET-OCre+ | P066_control | p66 | 0.1292 | 20.9 | 0.00618182 |
| P066_pkd1 | Pkd1^neo;Pax8+;TET-OCre+ | P066_PKD | p66 | 0.1509 | 19.8 | 0.00762121 |
| P066_pkd2 | Pkd1^neo;Pax8+;TET-OCre+ | P066_PKD | p66 | 0.1605 | 21.0 | 0.00764286 |
| P100_cont1 | Pkd1^neo;Pax8+;TET-OCre- | P100_control | P100 | 0.1427 | 27.0 | 0.00528519 |
| P100_cont2 | Pkd1^neo;Pax8+;TET-OCre- | P100_control | P100 | 0.1682 | 24.35 | 0.0069076 |
| P100_cont3 | Pkd1^neo;Pax8+;TET-OCre- | P100_control | P100 | 0.1636 | 24.1 | 0.00678838 |
| P100_pkd1 | Pkd1^neo;Pax8+;TET-OCre+ | P100_PKD | P100 | 0.2699 | 26.48 | 0.0101926 |
| P100_pkd2 | Pkd1^neo;Pax8+;TET-OCre+ | P100_PKD | P100 | 0.2726 | 23.7 | 0.01150211 |
| P100_pkd3 | Pkd1^neo;Pax8+;TET-OCre+ | P100_PKD | P100 | 0.2851 | 24.12 | 0.01182007 |
| P130_cont1 | Pkd1^neo;Pax8+;TET-OCre- | P130_control | P130 | 0.1719 | 26.13 | 0.00657865 |
| P130_cont2 | Pkd1^neo;Pax8+;TET-OCre- | P130_control | P130 | 0.1855 | 26.24 | 0.00706936 |
| P130_cont3 | Pkd1^neo;Pax8+;TET-OCre- | P130_control | P130 | 0.1766 | 28.26 | 0.00624912 |
| P130_pkd1 | Pkd1^neo;Pax8;TET-OCre+ | P130_PKD | P130 | 0.4353 | 28.57 | 0.01523626 |
| P130_pkd2 | Pkd1^neo;Pax8;TET-OCre+ | P130_PKD | P130 | 0.4801 | 24.91 | 0.01927338 |
| P130_pkd3 | Pkd1^neo;Pax8;TET-OCre+ | P130_PKD | P130 | 0.2875 | 25.31 | 0.01135915 |

**Table S1.** Information of the PKD and control mice used in this study. KW, kidney weight (g), BW, body weight (g).

| ID | ATAC reads per cell | RNA reads per cell | Total ATAC reads | Total RNA reads |
|---|---|---|---|---|
| P066_cont1 | 18039 | 1790 | 434726514 | 251698529 |
| P066_cont2 | 18609 | 2151 | 373422586 | 314163173 |
| P066_pkd1 | 16841 | 1941 | 375567214 | 299757192 |
| P066_pkd2 | 18181 | 1938 | 384306316 | 335314017 |
| P100_cont1 | 14042 | 2069 | 367700791 | 437579021 |
| P100_cont2 | 14922 | 1842 | 320085058 | 266088813 |
| P100_cont3 | 16300 | 2119 | 376577515 | 297138245 |
| P100_pkd1 | 15495 | 1969 | 346519417 | 324081795 |
| P100_pkd2 | 16846 | 2117 | 350093687 | 283236513 |
| P100_pkd3 | 18804 | 2354 | 323691557 | 276065421 |
| P130_cont1 | 17430 | 2092 | 346878407 | 339256963 |
| P130_cont2 | 6086 | 960 | 354380355 | 379407672 |
| P130_cont3 | 13308 | 1833 | 326734117 | 310507359 |
| P130_pkd1 | 15007 | 1988 | 346395931 | 284517476 |
| P130_pkd2 | 16605 | 2209 | 341717710 | 315837557 |
| P130_pkd3 | 9675 | 1836 | 324039235 | 326906052 |

**Table S2.** The numbers of total reads and reads per nucleus in the CellRanger output. ATAC reads per cell, ATAC Median high-quality fragments per cell; RNA reads per cell, GEX Median genes per cell; Total ATAC reads, Total sequenced read pairs for snATAC-seq; Total RNA reads, Total sequenced read pairs for snRNA-seq,

| Celltype | P66 | | P100 | | P130 | |
|---|---|---|---|---|---|---|
| | control | PKD | control | PKD | control | PKD |
| PTS1 | 21.4% | 22.6% | 19.3% | 23.2% | 25.2% | 22.8% |
| PTS2 | 22.7% | 22.8% | 21.8% | 25.8% | 21.1% | 22.5% |
| PTS3 | 5.5% | 5.2% | 5.7% | 4.2% | 5.1% | 4.0% |
| FRPTC_PEC | 1.1% | 2.1% | 0.5% | 2.7% | 0.4% | 5.5% |
| DTL1 | 1.3% | 2.1% | 1.9% | 1.5% | 1.7% | 1.7% |
| DTL2 | 0.2% | 0.5% | 0.3% | 0.4% | 0.4% | 0.6% |
| ATL | 0.6% | 1.1% | 0.9% | 0.8% | 0.9% | 0.7% |
| TAL | 12.2% | 14.8% | 13.6% | 12.5% | 10.7% | 11.3% |
| DCT | 6.5% | 3.4% | 6.8% | 4.0% | 5.8% | 4.8% |
| CNT | 4.0% | 2.7% | 3.9% | 2.2% | 4.8% | 3.3% |
| PC1 | 3.3% | 3.4% | 4.3% | 3.1% | 2.9% | 3.2% |
| PC2 | 1.0% | 1.0% | 1.4% | 0.8% | 1.1% | 0.9% |
| URO | 0.2% | 0.2% | 0.3% | 0.3% | 0.4% | 0.5% |
| ICA | 1.8% | 1.4% | 1.2% | 0.9% | 1.5% | 0.9% |
| ICB | 1.6% | 1.3% | 1.3% | 1.0% | 2.1% | 1.0% |
| PODO | 0.3% | 0.4% | 0.2% | 0.4% | 0.2% | 0.3% |
| ENDO | 12.7% | 12.0% | 11.9% | 12.2% | 9.2% | 11.1% |
| FIB | 1.8% | 1.1% | 2.7% | 1.1% | 4.2% | 1.4% |
| Myel | 1.1% | 1.1% | 1.2% | 1.8% | 1.7% | 1.6% |
| Bcell | 0.1% | 0.2% | 0.2% | 0.4% | 0.1% | 0.2% |
| Tcell | 0.4% | 0.4% | 0.3% | 0.6% | 0.5% | 0.6% |
| FAT | 0.1% | 0.1% | 0.1% | 0.2% | 0.1% | 0.2% |

**Table S3.** The frequency of nuclei for each cell type in the dataset of each time point / genotype.

| Genotype | Forward primer | Reverse primer |
|---|---|---|
| *Pkd1* flox allele | CCTGCCTTGCTCTACTTTCC | AGGGCTTTTCTTGCTGGTCT |
| *Pax8*$^{rtTA}$ | CCATGTCTAGACTGGACAAGA | CTCCAGGCCACATATGATTAG |
| Cre | AGGTTCGTTCACTCATGGA | TCGACCAG TTTAGTTACCC |

**Table S4.** Genotyping primers.
